# Multiscale Rheology of Aging Cancer Spheroids

**DOI:** 10.1101/2023.07.31.550652

**Authors:** Kajangi Gnanachandran, Massimiliano Berardi, Asmus Skar, Grażyna Pyka-Fościak, Joanna Pabijan, Javier Lopez Alonso, B. Imran Akca, Małgorzata Lekka

## Abstract

Cancer spheroids offer a valuable experimental model that mimics the complexity and heterogeneity of solid tumors. Characterizing their mechanical response is crucial for understanding tumor development, progression, and drug response. Currently, whole live spheroids are analyzed primarily using image analysis, which is challenging, requires extended incubation times, and has limited imaging depth. Here, we present a new label-free approach for characterizing sub-superficial structures of bladder cancer spheroids and measuring their mechanical response at three distinct stages of cancer progression. We study the microrheological changes induced by aging at the cellular and cluster levels by conducting a multi-physics characterization and modeling approach. We find that spheroids exhibit viscoelastic behavior that can be described by fractional models. We show that spheroids are mechanically heterogeneous, with strong depth and time-dependent variations associated with evolving structural features. Our approach opens new possibilities to study 3D *in vitro* models, paving the way for the discovery of novel and more precise procedure in cancer diagnosis based on the use of mechanomarkers.

## 1 Introduction

The field of mechanobiology has garnered increasing attention in recent years. The quantification of the viscoelastic properties of cells and tissues has helped understand cellular fate [1], tissue formation [2], and provided ways to distinguish between healthy and diseased cells [3]. Mechanomarkers have also shown promise in studying tissue-level pathologies [4, 5]: for example, nanoindentation devices used for breast tumor screening have reached the stage of clinical trials [6]. The majority of studies concerning mechanobiology have been conducted at either the single cell or tissue level, using various methods including atomic force microscopy (AFM), micro-indentation, optical tweezers, magnetic twisting, optical stretching, micropipette aspiration, acoustic radiation and scattering and Brillouin microscopy [3]. Although the usage of spheroids has considerably increased in recent years for studying cell-cell and cell-extracellular matrix (ECM) interactions in 3D, as well as the interplay between cells, ECM mechanics and drug diffusion and effectiveness [7–10], they are only partially explored in this field.

As pointed out by two recent reviews [11, 12], the current tools available to mechanobiologists can be split into surface methods, such as AFM [13], and bulk methods, such as parallel plate compression [14]. Considering how in these 3D models cell activity, metabolism, protein expression and drug response are tied to gradients of nutrients, pO_2_, pH, and waste, it is necessary to develop a method that can bridge current experimental approaches.

Currently, whole live spheroids are analyzed primarily using image analysis, relying on brightfield images for parameters such as size and circularity, and fluorescence for viability and internal structure [15]. However, image analysis is substantially more challenging in 3D cultures than in 2D cultures, because structures are observed from only one direction in brightfield imaging. Furthermore, extended incubation times are required for dye penetration, which only permits imaging to a depth less than the spheroid radius [16]. Tissue clearing, as well as more advanced microscopy methods such as light-sheet or multiphoton are potential solutions to the latter problem; however they represent either end point measurements or require advanced equipment [16]. According to a recent study on a large sample of spheroids, measuring the outer radius of spheroids is insufficient to predict necrotic/inhibited areas, and the most insightful information comes from the evaluation of the internal structures [17].

Here, we present microrheological studies on spheroids derived from bladder cells originating from different stages of cancer progression. We examine them at both the cellular and local cluster levels, and at two distinct time points during their growth. We couple the results obtained with AFM and Hydraulic Force Spectroscopy (HFS) [18] with the histological and confocal analysis of the spheroid inner structures. We present a new multiphysics experimental framework for HFS, coupled with a mathematical model that can retrieve depth-resolved mechanical properties of layered structures and infer structural changes, whilst minimizing sample preparation. We show how time-dependent remodeling of spheroids and depth-resolved mechanical response are paired, how spheroids exhibit power law rheology at multiple scales, and how those properties are associated with cell motility in three dimensions. Lastly, we describe how this approach could be implemented on a broader scale, leveraging its label-free nature and speed of execution.

## 2 Results

In this study, we prepared multicellular spheroids using three bladder cancer cell lines derived from distinct tumor stages, namely non-malignant HCV29 cells obtained from the non-cancerous part of the bladder carcinoma featuring the morphology and properties of the normal epithelium; HT1376 cells (grade III bladder carcinoma), and T24 cells (grade III transitional cell carcinoma). Spheroids are known to have a growth pattern similar to the one of solid tumors showing an initial phase of logarithmic character followed by a plateau [15]. We observed the evolution and the shape of the three spheroids over the course of their culture for two weeks. Figure 1-a shows the variation of the spheroid projected area over time. HCV29 and T24 spheroids shrink over time, reaching a stable plateau after about a week. In contrast, HT1376 spheroids kept on growing at a constant rate. All three spheroid types exhibited a high level of circularity, with a median value of ≈0.9. Circularity only showed minor variations between 3-day-old and 14-day-old spheroids (Figure 1-b), which were significant only in the case of HCV29 (p<0.01, Mann-Whitney U test). As cell-cell interactions, primarily governed by cadherins [19, 20], are important for the spheroid formation, we verified the expression of N/E cadherins using the Western blot technique (Figure 1-c). We found that HCV29 and T24 spheroids express N-cadherin and not E-cadherin, whilst the opposite is true for HT1376 spheroids, in agreement with previous studies done on 2D cell cultures [21]. Additionally, unlike the other two cell lines, HT1376 cells are known to be devoid of thick actin bundles [13], implying a possible role of the actin cytoskeleton organization in the development of the spheroids formed by them. We suggest that these could be the reasons behind the different behavior of the spheroids formed by HT1376 cells.

**Fig. 1.**
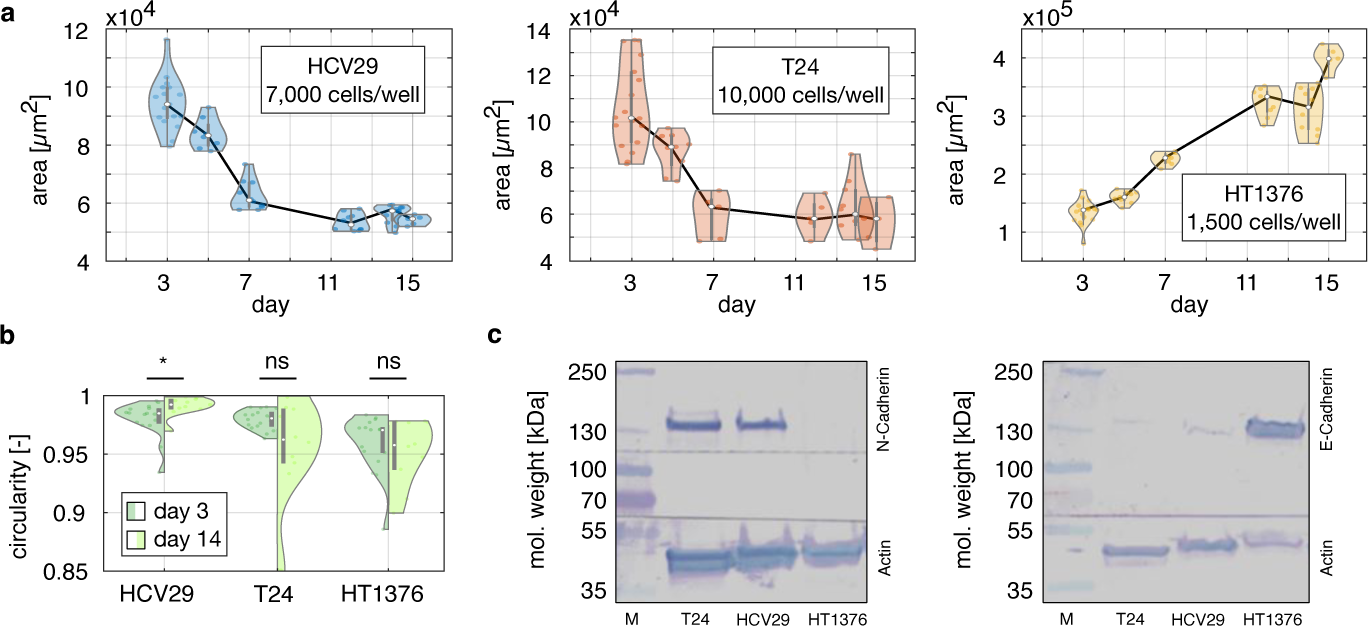
Results of the spheroid culture time in 96 U-shaped wells. A) Projected area over time, as measured from images recorded by an optical microscope. B) Circularity was determined for the bladder cancer spheroids on days 3 and 14. Violin plots represent the circularity of N≥10 spheroids. Only HCV29 spheroids show a significant change in circularity over the two weeks (tested with Mann-Whitney U test), with \**p <*0.01. C) Exemplary Western Blot results obtained for 3-day-old spheroids. HCV29 and T24 spheroids express only N-cadherin, whilst HT1376 spheroids expresses only E-cadherin. M stands for molecular weight standard (marker). The complete picture of the blots is reported in the supplementary figure S1.

### 2.1 Rheology of Spheroids is Time and Scale Dependent

We performed dynamic mechanical analysis (DMA) using both AFM and HFS, to characterize the spheroids’ rheological response at the cellular and cluster levels (Figure 2-a/b). The measurements were conducted after 3 days and 14 days of culture to evaluate the age-related changes in the biophysical properties of spheroids. In the case of AFM data, we modeled the frequency behavior using a standard microrheology approach: as reported by several researchers [22–24], cells tend to behave as power law materials when subject to dynamic nanoindentation. For this reason, we adopted a lumped parameter model composed of fractional elements. The repeating unit, named springpot [24], takes the form:

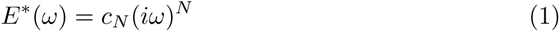

**Fig. 2.**
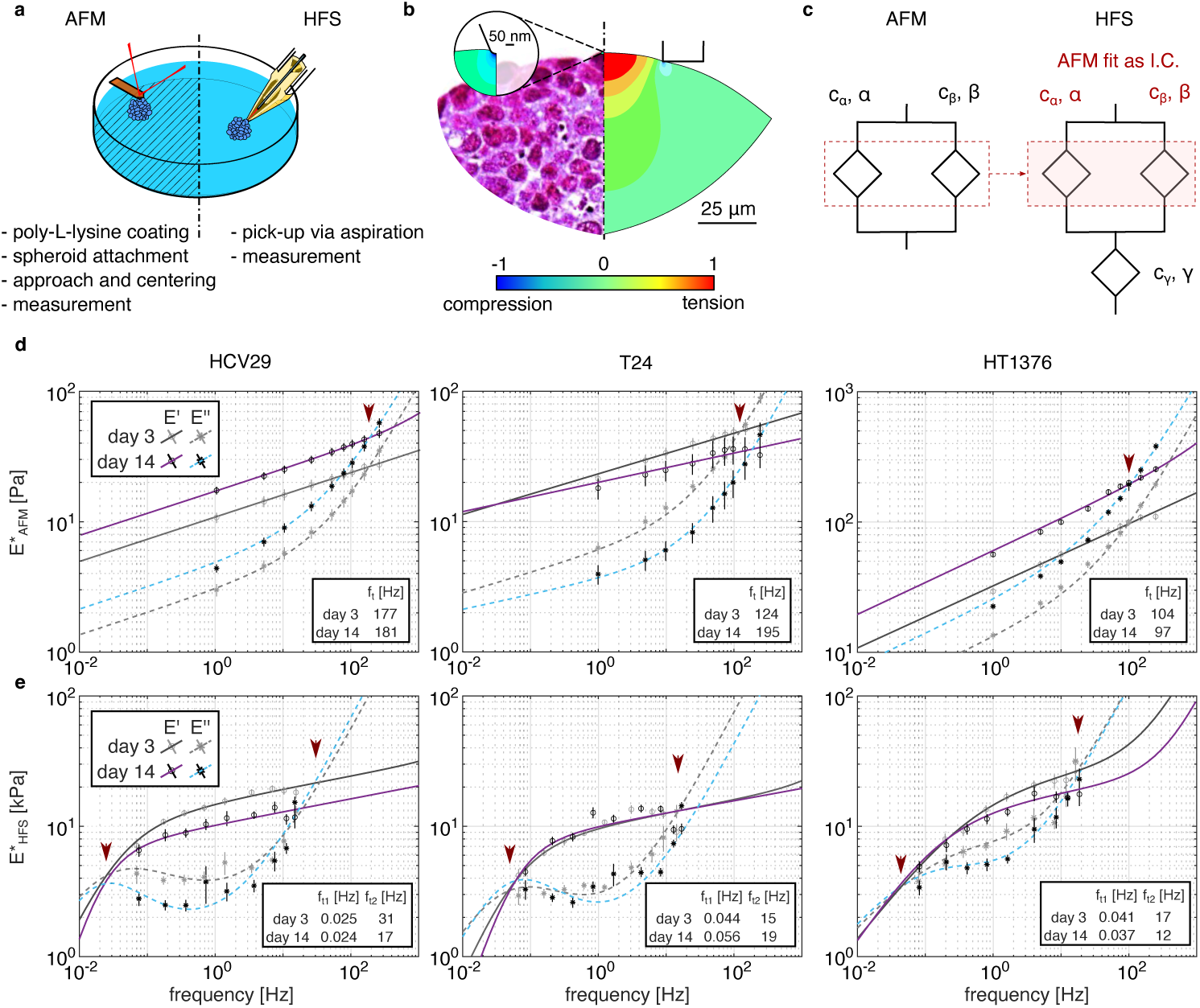
Spheroid microrheology at the cell (AFM) and cluster (HFS) scales and their evolution in time. A) Schematic of the workflow for the AFM and HFS approaches. B) Normalized displacement vector sum from FE models, showing the scale of indentation in spheroids (left - a histologically labeled fragment of the indented spheroid; right - FE simulation). C) Schematic representation of the lumped parameter models used. On the left, the equivalent of a double power law used to fit AFM data, and on the right the Poynting-Thompson model, used to fit HFS data. D) Microrheology results at both cell (AFM) and cluster (HFS) scales. Solid and dashed lines indicate storage and loss moduli, respectively. Grey: day 3; Blue and Purple: day 14. The red arrows indicate the transition regions, from a solid-like to a fluid-like regime, whilst the inset tables list the transition frequencies as estimated by the models. For each spheroid type, we performed n*_HF S_* =39-46 measurements on N=10-12 different spheroids, and n*_AF M_* =280-350 measurements on N=10-13 spheroids. Measured storage (circles) and loss (asterisks) moduli are shown as mean ± SEM, calculated by pooling all results together.

where *E^∗^* is the complex modulus in Pa, *c_N_* is a multiplying factor in Pa·s*^−N^*, *i* is the imaginary unit, *ω* is the excitation frequency in Hz, and *N* is the degree of derivation, between 0 and 1, with each springpot labeled with progressive lettering *α*, *β*. We modeled the AFM data with two springpots in parallel. A schematic depiction is shown in Figure 2-c (equations are presented in Methods). The model faithfully describes the material response over the three decades covered, for all lines and at all time points, as shown in Figure 2-d (first row). All three spheroid types exhibit a solid-liquid transition, located at 177 and 181 Hz (for HCV29 spheroids), 124 and 195 Hz (for T24 spheroids), and 97 and 104 Hz (for HT1376 spheroids). HCV29 and HT1376 spheroids stiffened over two weeks, keeping their transition frequency and loss factor (defined as *E^′′^/E*) values constant, whilst T24 spheroids show a weaker *E^∗^* dependency on frequency after two weeks, and as such, sees the transition frequency increase considerably, to almost twice the initial value. Unlike the other two types of spheroids, in T24 spheroids the loss factor is reduced by about 30%, approximately evenly across the tested range.

Looking at the numerical results of the fit (Table 1), it is possible to appreciate how the values obtained align with prior reports on cell mechanics, particularly in terms of fractional derivative order [13, 22]. It is worth noting how the *β* term is generally close to 1, indicating purely dissipative behavior. At the same time, *c_β_* has a rather large error in magnitude, comparable to its optimal value (see Table 1). One possible explanation is that the model does not account for additional dissipative events occurring in the spheroids. Other possible reasons could be dissipation caused by cell-cell interactions and poroelastic relaxation; however, given the scale of the test, the diffusivity of biological materials, and the frequency range probe, the latter is likely a limited effect [18]. Cell-cell interactions certainly contribute to the response measured by AFM, however, considering the volume subject to deformation (see Figure 2-b for a qualitative depiction), the contribution of interfaces is too small to be noticeable. This contrasts with semi-local techniques, such as micropipette aspiration, as schematically shown in Figure 2-b.

**Table 1.** Results of the parallel springpots model (AFM) and Poynting-Thompson model (HFS) at 3 and 14 days, shown as mean ± standard deviation.

| condition | $c_\alpha$ [Pa·s$^\alpha$] | $\alpha$ [-] | $c_\beta$ [Pa·s$^\beta$] | $\beta$ [-] | $c_\gamma$ [Pa·s$^\gamma$] | $\gamma$ [-] |
| --- | --- | --- | --- | --- | --- | --- |
| HCV29 spheroids |  |  |  |  |  |  |
| AFM (3) | 11.36 ± 2.03 | 0.17 ± 0.07 | 0.11 ± 0.14 | 1.00 ± 0.20 | - | - |
| HFS (3) | 15.65 ± 2.90 | 0.10 ± 0.16 | 0.53 ± 1.22 | 1.00 ± 0.60 | 199 ± 550 | 0.83 ± 0.73 |
| AFM (14) | 17.89 ± 2.65 | 0.17 ± 0.06 | 0.23 ± 0.26 | 0.95 ± 0.17 | - | - |
| HFS (14) | 10.42 ± 1.57 | 0.10 ± 0.18 | 0.66 ± 1.11 | 1.00 ± 0.45 | 367 ± 1851 | 1.00 ± 1.47 |
| T24 spheroids |  |  |  |  |  |  |
| AFM (3) | 23.87 ± 4.33 | 0.16 ± 0.07 | 0.30 ± 0.28 | 1.00 ± 0.15 | - | - |
| HFS (3) | 10.35 ± 2.53 | 0.10 ± 0.21 | 0.71 ± 1.20 | 1.00 ± 0.43 | 106 ± 289 | 0.74 ± 0.86 |
| AFM (14) | 20.30 ± 6.98 | 0.11 ± 0.13 | 0.15 ± 0.37 | 1.00 ± 0.30 | - | - |
| HFS (14) | 10.04 ± 1.27 | 0.10 ± 0.13 | 0.58 ± 0.74 | 1.00 ± 0.33 | 163 ± 329 | 1.00 ± 0.62 |
| HT1376 spheroids |  |  |  |  |  |  |
| AFM (3) | 65.06 ± 12.87 | 0.24 ± 0.07 | 1.45 ± 1.35 | 0.95 ± 0.13 | - | - |
| HFS (3) | 23.24 ± 73.24 | 0.10 ± 1.00 | 1.59 ± 11.38 | 0.97 ± 1.20 | 43 ± 260 | 0.63 ± 1.40 |
| AFM (14) | 34.74 ± 5.50 | 0.24 ± 0.60 | 0.60 ± 0.55 | 1.00 ± 0.14 | - | - |
| HFS (14) | 16.94 ± 17.90 | 0.10 ± 0.47 | 1.35 ± 4.02 | 1 ± 0.51 | 33.59 ± 84.26 | 0.61 ± 0.55 |

We repeated the DMA with HFS to probe the response of spheroids at the cluster scale. The results (Figure 2-e) show a vastly different behavior compared those collected by AFM, with the storage modulus almost plateauing towards the higher frequency range, the loss modulus dominating at low frequencies, and the presence of two transition frequencies: a liquid-solid transition at low frequency, in the scale of tens of seconds, and a solid-liquid transition at high frequencies, in the scale of tenths of seconds. Some feature of cellular mechanical response seem apparent also in the HFS measurement, such as the steeper frequency dependence on the HT1376 spheroid response and the increase in transition frequency over time for T24 spheroids. Overall, the response of spheroids has a strong viscoelastic character at multiple scales, and is also dependent on spheroid age.

### 2.2 Classical Lumped Parameter Models are Inadequate for Describing Mesoscale Spheroid Rheology

The behavior measured by HFS, particularly at low frequency, is akin to previous observations on spheroid mechanics, which showed them behaving like liquids upon applied suction pressure. We first fit our frequency data to a common lumped parameter model used for spheroids, i.e. a Burgers model, which is also known as the modified Maxwell model [14, 25–27]. This model consists of a standard linear solid model in series with an additional dashpot. In all cases we could find in the literature, its parameters were determined by fitting experimental data in time domain either by using the Boltzmann hereditary integral approach [25] or following a step response [14]. Interestingly, we found this model to be completely inadequate for capturing both phase transitions. The results of the fitting procedure using classic lumped parameter models are available in the Supplementary material, whilst the methodology is detailed in the Methods section. Previous results used experimental data primarily obtained through video tracking at relatively low frequency. The higher temporal resolution arising from the interferometric readout makes it clear that the rheological behavior of spheroids cannot be described by a limited number of elements with single relaxation times.

To model this response, we once again resorted to fractional models [24]. We hypothesized the material behavior apparent from the HFS measurement could be crudely modeled by putting a “cell element” in series with an “interface element”, which in this context describes cell-cell (mostly mediated by cadherins) and cell-ECM interactions. We put a single springpot in series with the cell model for simplicity and to be in line with the Burgers models. This is known as the Poynting-Thompson model [24], and is shown schematically in Figure 2-c (for the equation, see Methods).

We initialized the parameters of the parallel terms with the results obtained from AFM microrheology to guide the fit. Despite its simplicity, we found that this approach could accurately capture the spheroid response, in particular predicting both low and high frequency transitions. This is true regardless of the spheroid type and time point. In several cases, the optimal parameters of the parallel springpots were within 2*σ* of the AFM estimate (see Table 1). In all conditions, the additional springpot had a *γ* close to one, indicating predominantly dissipating behavior at the interfaces. This model suggests that the rheological fingerprint of a cell line is measurable even when the scale of testing is significantly larger. At the same time, this approach implies that if the cell mechanical properties and their packing conditions are known, the interfacial properties can be estimated.

### 2.3 Low Frequency Transitions are correlated to Spheroid Fusion Dynamics

Our model predicts a very low transition frequency. As the time scale is comparable to that of collective cell movement [28], we hypothesized this could be linked to unjamming transition and cell motility within the spheroid [29]. To verify this hypothesis, we conducted homotypical spheroid fusion experiments (Figure 3), measuring the degree of spheroid fusion after 24 hours of first contact and relating it to the lower liquid-solid transition frequency found with the Poynting-Thompson model (defined as *Re*(*E^∗^*) = *Im*(*E^∗^*)).

**Fig. 3.**
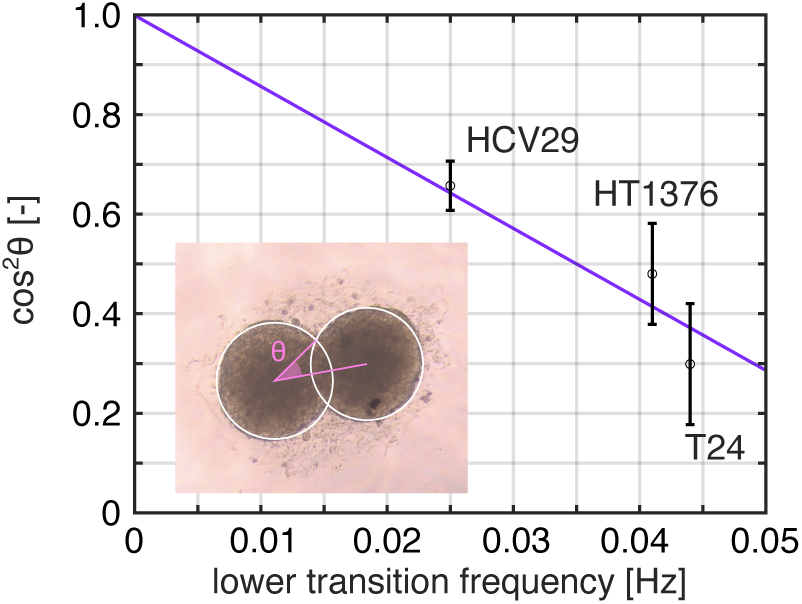
Relationship between the squared fusion angle 24 h after fusion initiation and the lower transition frequency identified by the Poynting-Thompson fit on HFS data. In the inset, an exemplary image (HT1376 spheroids fusion) shows how sin*θ* is calculated.

Interestingly, we found a monotonic relationship between the quantities, as shown in Figure 3. Whilst this result should be taken with care, as the lower transition frequency is a mathematical extrapolation rather than a directly measured quantity, this result corroborates the hypothesis of the series term roughly describing interfacial phenomena and is in agreement with other studies [29, 30]. The line in the graph results from a linear fit of mean values (R^2^=0.86, slope=-14.32 s), where we imposed an intersect of (0,1), as the limit behavior for which two solids would simply never fuse.

### 2.4 Spheroids Undergo Time and Layer Dependent Structural Remodeling

A limitation of the lumped parameter model description based on DMA is that it does not consider the actual stress dissipation across the elements, nor their spatial distribution. These factors are crucial for the proper quantification of micromechanical behavior in composite systems [31]. Moreover, the lumped parameter model cannot explain why the spheroids appear softer after 14 days when tested with HFS, in antithesis with the AFM data, nor what process is causing the change in transition frequencies. On top of that, the high uncertainty of some parameters suggests that the material cannot be fully described under the assumption of isotropicity and homogeneity.

To motivate these changes, we investigated the inner structure of the spheroids by histological and confocal analyses. Histological analysis (Figure 4-a) of the three types of spheroids over time shows a layered structure with a denser, highly proliferating outer layer. At day 3, all spheroids feature proliferating cells in the inner region as well. In addition, HT1376 spheroids display vacuoles, which may be a precursor to the necrotic core establishment. At 14 days, T24 spheroids show denser packing of cells across the section. HCV29 spheroids experience a similar but milder increase in the outer regions. HT1376 spheroids, on the other hand, have a necrotic core as a result of continuous growth and limited nutrient penetration. The necrotic area is off-center, likely because of the culture method. Masson’s Trichrome and Sirius red stains highlight the presence of small quantities of collagen in T24 spheroids at day 3, which is not present in spheroids after 14 days of culture. On the other hand, HCV29 spheroids show a considerable amount of collagen deposition, especially in the inner region.

**Fig. 4.**
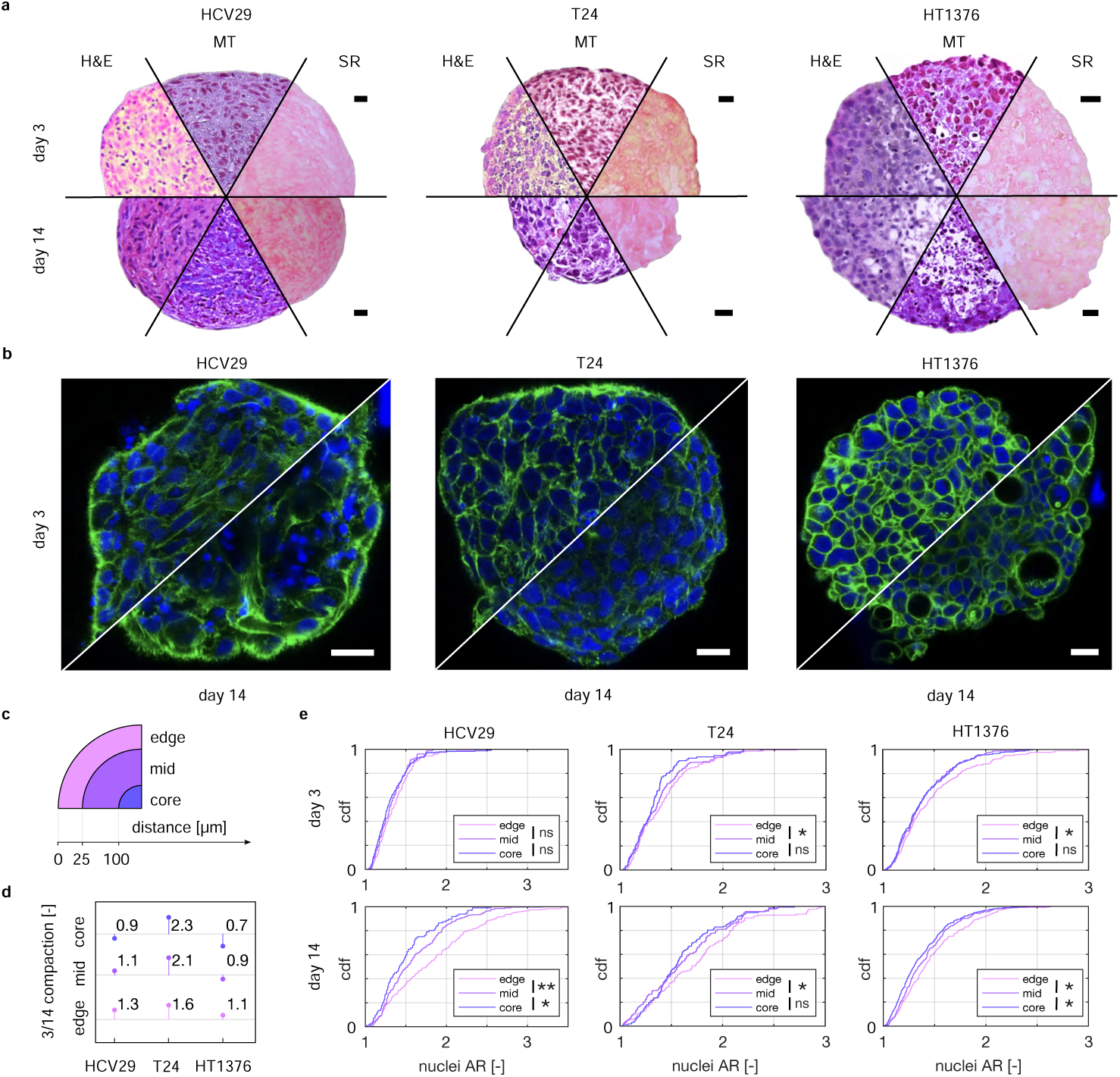
Structural evolution of the three spheroid types over time. A) Histological staining of the spheroids: Hematoxylin & Eosin (H&E), Masson Trichrome (MT), and Sirius Red (SR). Spheroid cross-sections at two time culture points are shown. B) Confocal images showing the organization of actin cytoskeleton inside spheroids. Green: F-actin, stained with phalloidin conjugated with Alexa Fluor 488, blue: nuclei, stained with Hoechst 33342. C) Schematic subdivision of the spheroids in three regions: edge, mid, and core, as a function of the distance from the surface. D) Compaction over time, per layer and by line, measured as the ratio between the number of nuclei per *µ*m^2^ at 14 over 3 days. E) Empirical cumulative distribution functions (cdf) showing the aspect ratio distribution of the nuclei, categorized in three characteristic zones inside the spheroids. Significance based on Mann-Whitney U test, with \**p <*0.05, \*\**p <*0.0005. All scale bars are 20 *µ*m. The extended versions of the histological staining and confocal images are illustrated in supplementary figures S2 and S3.

Confocal analysis (Figure 4-b) shows the evolution of actin organization in aging spheroids. At day 3, actin is mainly grouped around the cells, albeit it is possible to still see some actin fibers in HCV29 and T24 spheroids. After two weeks, HCV29 spheroids display an irregular actin cytoskeleton and condensed nuclei, a sign of apoptotic cells. In T24 and HT1376 spheroids a well-organized actin cytoskeleton and marginally smaller nuclei are present with respect to their younger counterparts. In the case of HT1376 spheroids, the vacuoles observed in the histology are also visible here.

Quantitative analysis of hematoxylin & eosin (H&E) histology images (Figure 4, c-e) shows how all three spheroid types undergo significant remodeling. An increase in nuclei density in the outer region over time and an increase in the nuclei aspect ratio (AR) were observed. The time-dependent trends are particularly marked in the lines expressing N-cadherins (median AR of the proliferating layer increases ≈50% for both HCV29 and T24 spheroids). The only exception (lowered cell density in the inner region of HT1376 spheroids) can be linked to the presence of a necrotic core. In all lines, after 14 days the outer layer featured a nuclei AR distribution significantly different from the inside (Mann-Whitney U test).

Taken together, these results give some insight into the nature of the discrepancies measured by DMA. As the AFM measurements are sensitive to the surface properties of the sample, the increase in cell density in the outer shell appears to be a good explanation. HCV29 spheroids see the cell packing density increase over time, as well as a small amount of collagen deposited in the proliferating zone. Mechanically, they stiffen while maintaining the same viscous character and transition. T24 spheroids lose the small amount of collagen displayed at day 3 over time, but at the same time experience a more severe increase in packing density, which may explain why the apparent stiffness remains unaltered with a reduction in viscous character. While HT1376 spheroids demonstrate minor density gain and no volume stabilization, they also stiffen. At the same time, the distribution of nuclei aspect ratios appears essentially identical to that of day 3, making an argument for cytoskeletal alignment-related stiffening unlikely. Stiffening with the increasing number of divisions has also been observed in other studies, especially for epithelial cell lines, even though the origin of this is unclear [32, 33]. The change in aspect ratio also provides a hint for the change in response measured via HFS, as the cell/cell surface in the radial direction changes considerably.

### 2.5 HFS Tomography Reveals Subsuperficial Mechanical Heterogeneities in Live Spheroids

The results presented so far clearly indicate that modeling spheroids under the assumption of homogeneity yields ambiguous results, and also appears insufficient in describing the layered nature of the samples. While there have been attempts to predict layered elasticity from nanoindentation data [34], the available probes are too small to explore spheroid mechanics across the layers we specified. Custom AFM probes are a potential solution in this regard; however, increasing size only increases errors associated with double contact, particularly in the case of asymmetric structures like the HT1376 spheroids at 14 days [35]. For these reasons, we extended the HFS method, to retrieve and model the full displacement profile of the sample across its diameter. For this purpose, we used the Adaptive Layered Viscoelastic Analysis (ALVA) software [36] (see Figure 3-a for a visual summary of the approach). In this framework, the spheroid is viewed as a collection of flat layers, each of finite thickness, resting on a semi-infinite medium. Because of the size of the pipette and its associated non-zero strain field, we limited the analysis to two layers, optimizing for both elastic moduli and top-layer thickness. A detailed procedure is described in the Methods section. At the same time, we measured spheroid cryosections with AFM, akin to what is commonly done in tissue mechanics[37]. The underlying assumption of these measurements is that the composition throughout the thickness (about 20 *µ*m) is homogeneous, and the surface has low roughness and low adhesion. We used Hertzian modeling to obtain a baseline for mechanical response to compare the tomography analysis. Doing so neglects the viscous component of the mechanical response. However, given that the topology of the springpots in both time and scale is consistent, this approach can still be used to make statements about relative changes.

These additional results, albeit limited to quasi-static properties, provide a much clearer picture of the mechanics of the spheroids under study. Figure 5-b shows an example of a displacement profile along the spheroid diameter, and the corresponding fit obtained with ALVA (top panel). Despite the approximation of purely elastic behavior and lack of curvature effects, a finite element simulation with the same parameters (E*_top_*/E*_mid_*=5.8, t*_shell_*=19.5 *µ*m, Figure 5-b, lower panel) gives a remarkably close result. Repeating the procedure on all suction ramps of DMA data shows how the spheroids under study tend to be composed of a stiff shell and a significantly softer core. This measure partially confirms what has been indirectly observed under parallel plate compression in a previous study [38]. However, closer inspection of the Young’s moduli ratio distribution (Figure 5-c), reveals a broader range of possible conditions. After two weeks, HCV29 spheroids have a median E*_shell_*/E*_mid_* of 1.55 (down from 2.62 for spheroids on day 3), with half the spheroids featuring at least a measure indicating a stiffer core. As this change is driven by a reduction of the modulus of the outer layer (p<0.003, whilst the change for the core is non significant), this could be explained by a different cell-cell versus cell-ECM adhesion, as the collagen deposition is predominant past the proliferative rim,as shown by the increased stiffness measured by AFM on cryosections (Figure 5-f). T24 spheroids feature a comparatively larger E ratio (median 5.07 at day 3, 3.75 at day 14), which is higher than the AFM estimates (1.21 and 1.04 respectively), but with the same tendency. The HFS measurements of HT1376 spheroids feature the highest ratio (10.01 for spheroids on day 3, 15.19 for spheroids on day 14). This result is particularly interesting when compared to AFM, as in the cryosection measurements the trend is not visible, with median values between 2 and 3 kPa in all regions and at all times. Whilst the increase can be associated with the presence of the necrotic core, whose lack of mechanical contribution under tension is not perceived in cryosections, the difference in elastic moduli is already very strong at day 3 (median of 10) prior to its establishment. We believe two concurrent explanations are likely: on one hand, the presence of vacuoles at day 3 reduces the resisting cross-section of the internal structure, and on the other hand the internal hypoxic environment could be associated with the attenuated expression of E-cadherin, as reported in previous works [39, 40].

**Fig. 5.**
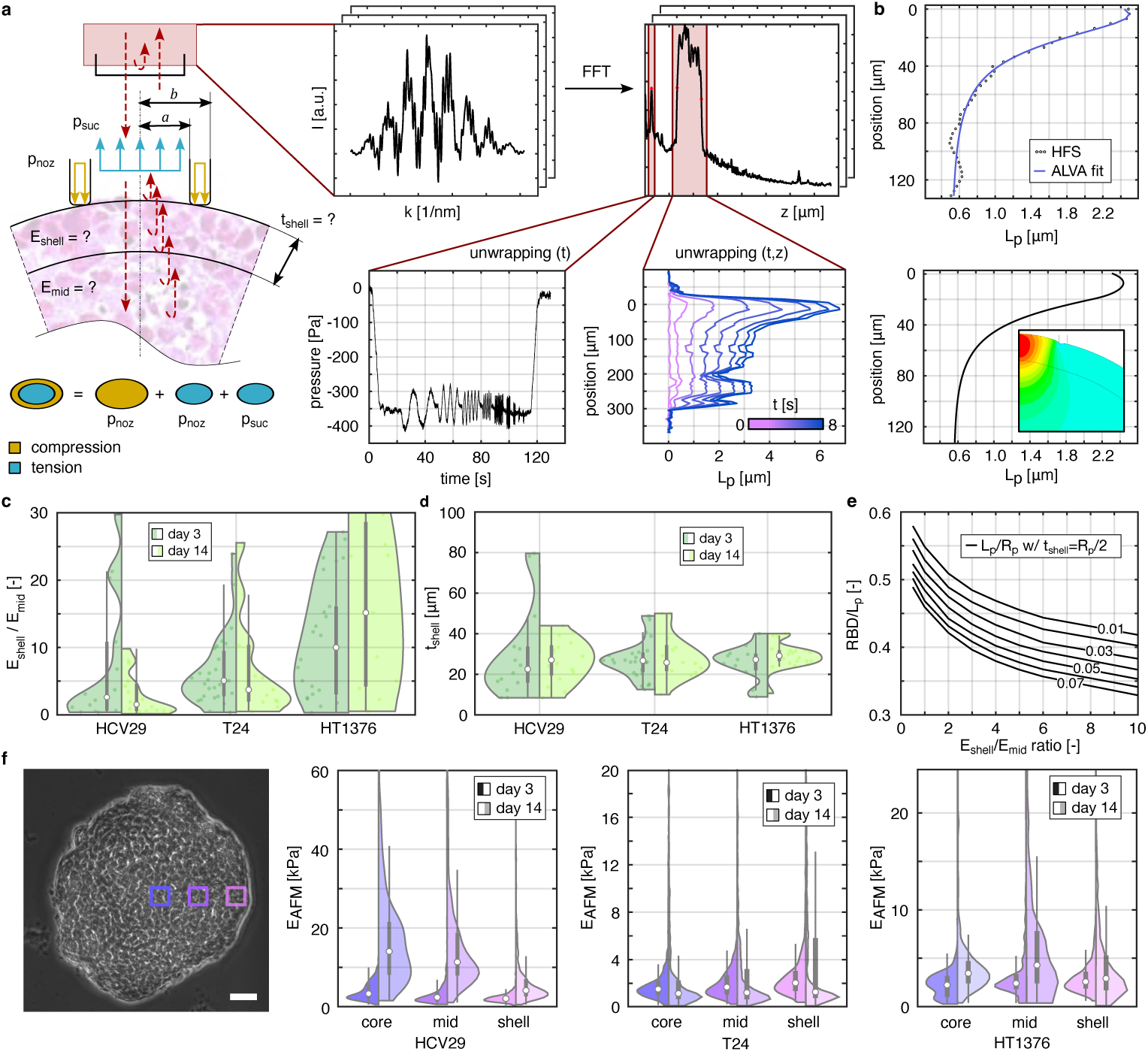
Depth-resolved mechanical properties of cancer spheroids. A) Schematic workflow of spectra acquisition, isolation, and modeling with ALVA. In the bottom right graph it is possible to see how the spheroid deforms within the nozzle while rigidly displacing at the distal end. B) In the top panel, experimental data (T24 spheroids, 14 day) vs ALVA estimate. In the bottom panel, the results of a FE simulation initialized with the parameters obtained with ALVA. C) Distribution of E*_shell_*/E*_mid_*of the layered elastic fit. The violin plots show pooled results from N≥10 with at least 10 spheroids measured D) Distribution of the estimated thicknesses of each layered elastic analysis. The violin plots show pooled results from N≥10 with at least 10 spheroids measured E) Results of a series of FE simulations showing how the ratio between aspirate length and rigid body displacement is tied to the relative layer properties. F) Results of the AFM analysis on cryosections. The example phase contrast image belongs to a HCV29 spheroid at day 3. The scale bar is 50 *µ*m. The violin plots show N≥300 pooled results from 3 independent slices for each condition.

Notably, despite the broad range of relative moduli, the estimated shell thickness is rather consistent in time and across lines, with a median of around 26 *µ*m and an interquartile range between 18 and 35 *µ*m (see Figure 5-d). The values are comparable to the thickness of the proliferative rim that we evaluated qualitatively from the histological analysis (Figure 4, panels a and c-e), two to three cells thick. The consistency of this result is useful because it implies that our findings could be used, to a degree, also in micropipette aspiration measurements using relying on image analysis. We simulated a few aspiration experiments, assuming a shell thickness of 25 *µ*m and an aspirating pipette of 50 *µ*m in diameter, varying the stiffness ratio of the shell and internal structure from simulation to simulation. We found that the ratio between rigid body displacement at the distal end over the aspirated length univocally identifies the E ratio of a 2-layered body. The results, shown in Figure 5-e, are an example of a look-up table that could be used for such a purpose.

AFM and HFS, based on quasi-static analysis, provide complementary information, highlighting the behavior of the cytoskeleton or fibrous ECM and interfacial properties respectively. More sophisticated HFS models could infer cell properties, as hinted by the DMA results. This result could guide the choice of a measurement technique based on the type of investigation. For example, HFS may be preferred when studying drug treatments that alter cadherin expression [41], whilst AFM may be more insightful when studying cytoskeletal-targeting drugs [42]. Altogether, these results appear to remove part of the ambiguity in mechanical response; nevertheless they come with a caveat: the interpretation of ratios, and their variations, was performed in light of structural information obtained via histological analysis. This means that the interpretation is still complicated if only mechanical tomography data is available.

### 2.6 Intensity Fluctuation Analysis of HFS Data Shows Unique Patterns for Different Internal Spheroid Structures

To mitigate the abovementioned issue, we implemented techniques commonly used in Optical Coherence Tomography [43, 44]. Specifically, we analyzed the HFS intensity profiles (figure 5-a) after capturing the sample, but before the actual mechanical test. We isolated the portion of the signal corresponding to the sample, akin to what we did in tomography, and measured its variance (LIV), as well as the speed of decorrelation (OCDS). For a detailed description of how these quantities have been calculated, see Methods. While these results lack the molecular specificity of, for instance, a confocal image, they can still provide a guide for investigations: lower LIV values indicate that the intensity signal has minimal fluctuation, i.e. there is less motion of scattering elements. The OCDS signal, given the time interval within which it is calculated, is sensitive to slow tissue dynamics [45].

The results, summarized in Figure 6, show interesting patterns. We reported the full cross-section results for completeness, but the results of the left half are affected by the capture pressure, as the samples partially behave as liquids. The following discussion refers to differences seen in the right half of the cross sections. All three spheroids show approximately constant LIV and OCDS on day 3. After two weeks, however, they feature different responses. Both the LIV and OCDS signals of HCV29 spheroids see a dip at the center of the spheroid, with the signal increasing towards the surface. This could be associated with the ECM deposition and reduced cell density in the inner area. The signals from T24 spheroids appear constant and comparable in magnitude to the data collected on day 3. HT1376 spheroids feature a dip in the OCDS signal, while LIV stays flat. This result is compatible with the presence of a necrotic nucleus and its consequential lack of dynamics.

**Fig. 6.**
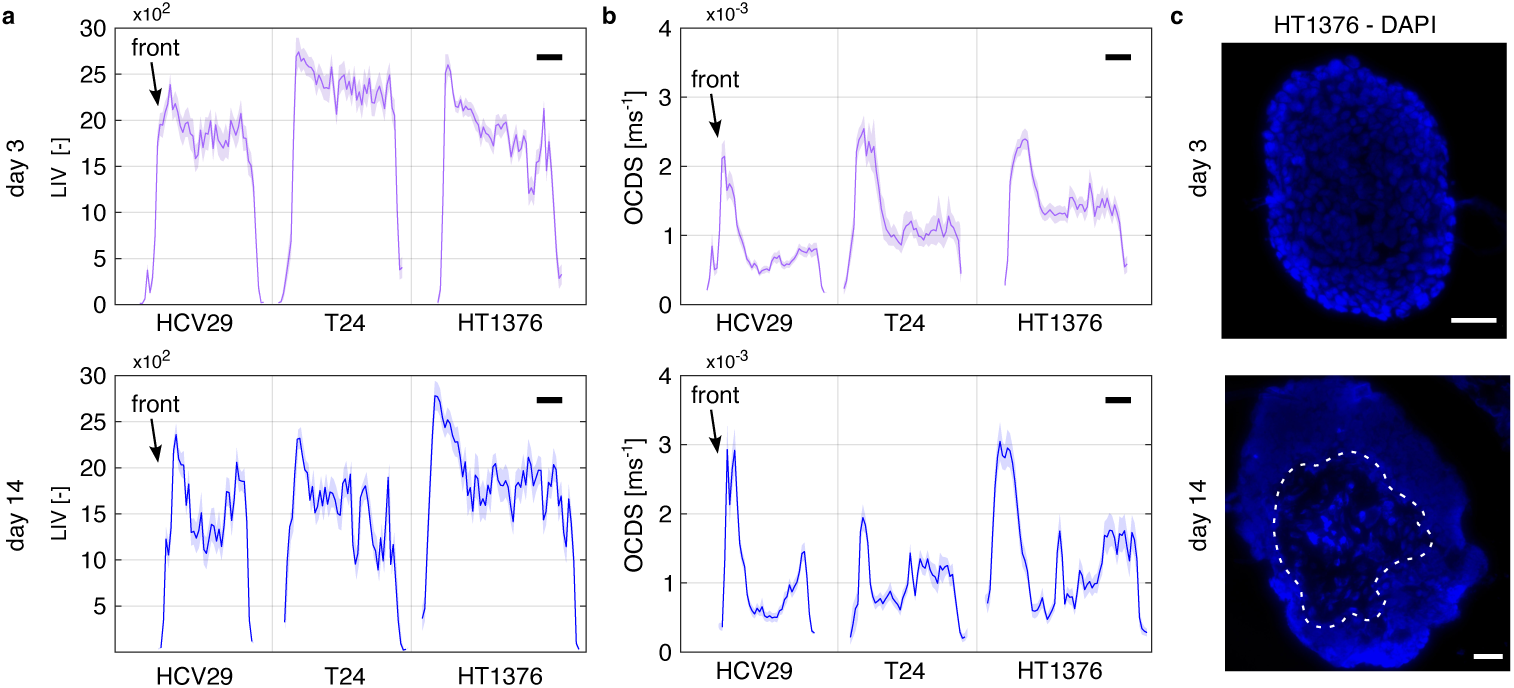
Results of the intensity fluctuation analysis of HFS signals. A) Results of the LIV calculations, at 3 (top) and 14 (bottom) days. B) Results of the Autocorrelation decay speed calculations at 3 (top) and 14(bottom) days. In all cases, the detection fiber was located on the left side, as denoted by the black arrows. In all cases, plots are displayed as mean ±SEM, with N=10-13 spheroids per condition. Scale bars are 50*µ*m. C) Fluorescence imaging (nuclei, stained with DAPI) of cryosections. The example shown belongs to HT1376 spheroids. In the bottom picture, the dashed white line denotes the necrotic region. All scale bars are 50 *µ*m. The exemplary images for all the other spheroids can be found in the supplementary figure S4.

## 3 Conclusions

The mechanical properties of bladder cancer cells have been extensively measured by AFM, showing that non-malignant cells are stiffer than cancer cells [46–48]. However, the majority of these studies were done at the single-cell level. In our recent study we showed that the overall elastic properties of cells have a similar tendency both as single cells and multicellular spheroids, and that rheological mechanomarkers are a better way to discriminate the cells deriving from various stages of bladder cancer progression [13]. Here, we took a step further and performed rheological measurements on cancer spheroids during their development at two different organization levels: single cells (AFM) and clusters (HFS). The experiments we presented in this work describe the age-related changes in the viscoelastic properties of cancer spheroids and they confirm the ability of the mechanomarkers to differentiate the non-malignant cells HCV29 spheroids from the more advanced tumor spheroids formed from T24 and HT1376. More importantly, we show how the mechanical properties of the spheroids formed by these different cell lines are correlated to their inner structure and the presence of ECM. Thus, we point out that treating spheroids like homogeneous bodies results in an incomplete depiction of their structure and mechanical response.

The information retrieved from the tomography data, both in terms of mechanics and autocorrelation information, proved particularly insightful, despite the lack of time-resolved information. It is however important to point out that this is possible down the line, as the displacement fields are resolved in time to less than a millisecond. Further studies could implement depth-resolved viscoelastic analysis and include the possibility of layer delamination, a feature already present in ALVA that wasn’t explored in this work. The first aspect is particularly crucial, as recent studies have shown how cellular response to the viscosity of its microenvironment outweighs rigidity sensing [49, 50].

The HFS technique is currently limited to performing sequential measurements, which limits its throughput. This is, however, mitigated by the fact that the current embodiment of the HFS method allows for targeting specific regions of a spheroid, which is not afforded in microfluidic approaches. Whilst the samples we studied showed a good degree of axial symmetry, multiple reports in the literature show how this is not the rule [51, 52]. The ability to target specific regions could also be useful to measure specific locations of otherwise hard-to-access samples, such as fusing spheroids. Thanks to its slenderness, the current pipette design allows in principle for measuring up to 1536 wells in a well plate format. This feature could be readily implemented by mounting the HFS pipette on, for example, a pipetting robot.

Given the usefulness of spheroid in modeling drug uptake in solid tumors [53–57], and considering how spheroid age affects diffusivity and toxicity [58], we believe our approach holds great potential for studying the interplay between mechanical response and drug action [42, 59, 60], as well as for studying the effects of ECM stiffening/softening and their influence on drug treatments [10, 61, 62].

## 4 Methods

### 4.1 Spheroid culture

We cultured spheroids from three cell lines: HCV29 – non-malignant transitional cancer of the ureter (established at the Institute of Experimental Therapy, PAN, Wroclaw, Poland) [63], T24 – transitional cell carcinoma from ATCC (LGC Standards, Poland) [64] and HT1376 – grade III urinary bladder cell carcinoma from ATCC (LGC Standards, Poland) [65]. HCV29 and T24 were cultured in the Roswell Park Memorial Institute Medium (RPMI) 1640 medium (Sigma-Aldrich, Poznań, Poland) supplemented with 10% of fetal bovine serum (FBS, Sigma-Aldrich, Poznań, Poland). HT1376 were grown in Eagle’s minimum essential medium (EMEM, LGC Standards, Poland) with 10% FBS. All cell lines were grown in culture flasks (Sarstedt, Germany) in an incubator (Nuaire, USA) at 37° C in 95% air and 5% CO2 atmosphere. We kept relative humidity above 98%, and carried out cell passaging at 80-90% of the confluency level. For HCV29 and T24 cells, 0.05% and for HT1376 cells, 0.25% of trypsin-EDTA solution (Sigma-Aldrich, Poznań, Poland) was applied. After a few passages (< 10), cells were ready to be seeded on 96-well U-bottom 3D cell culture plates (Thermofisher). Depending on the cell line, the number of cells/well was adjusted to obtain spheroids of 350-400 *µ*m diameter (for HCV29 spheroids: 7000 cells/well, for T24 spheroids - 10000 cells/well and for HT1376 spheroids- 1500 cells/well were used).

### 4.2 Spheroid fusion

For the spheroid fusion experiments, spheroids were prepared from the three types of bladder cell lines as described above. After 3 days of culture, two spheroids were put together in a new well with the corresponding fresh culture medium for each cell line. The degree of the fusion of the homotypical spheroids was determined after 24 hours starting from their first contact.

### 4.3 AFM microrheology

To immobilize the spheroids prior to indentation, we coated Petri dishes with 0.01% poly-L-lysine (Sigma-Aldrich, Poznań, Poland) and 5% glutaraldehyde (Sigma-Aldrich, Poznań, Poland). Briefly, we cleaned the Petri dish using ethanol and deionized water and dried it. The bottom Petri dish surface was incubated with diluted glutaraldehyde for 20 minutes and washed with water. Afterward, the diluted poly-L-lysine was placed for 30 minutes. We then remove the solution and let the dish dry. At this point, it was ready for use. We collected the spheroids from the culture plates and placed them on the coated dishes. Before starting a measurement, we waited 15-30 minutes to ensure that spheroids adhered to the poly-L-lysine-coated surface. The microrheological measurements were conducted using a NanoWizard IV (Bruker, JPK Instruments) in PBS and at room temperature. We used an NSC36 cantilever (0.6 N/m, tip height 12-18 *µ*m and half-open angle of 20 degrees). We set the approach velocity to 3 *µ*m/s, and the force setpoint to 30 nN. A high force set point enables to probe deeper regions of the spheroids. Since such a threshold may lead to membrane ruptures, we inspected every curve and analyzed only those that complied with the Hertzian response. An example of the recorded force curve is shown in the supplementary figure S6. The dynamic response was set between 1 and 250 Hz in 9 log-evenly spaced frequencies. The following parameters were set: a constant oscillation amplitude of 50 nm for 10 oscillations, sampling 600 points per period, and waiting 0.5 seconds between test frequencies. We collected the measurements in a series of matrices (9 *µ*m^2^, 2 maps/spheroid), on 12-15 spheroids per type and time point. We did not apply any hydrodynamic drag correction. We qualitatively inspected the need for it by observing the separation between approach and retract curves in the quasi static indentation at comparable velocities, and we found the baselines to be essentially overlapping. This may be due to the relatively high tip length, spheroid curvature, and soft substrate.

We analyzed the raw data in JPK Data Processing using the Hertzian contact model for a pyramidal indenter with blunted end assuming a conical shape (for details, see the Supplementary material). Then, we exported the results (storage, loss and corresponding frequency values) in *.csv files and imported them in Matlab R2022b to fit the full set of measurements per line/day in the complex plane to the equation:

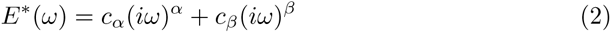

where *α* and *β* are the indices of derivation of the two springpots (between 0 and 1), and *c_α_* and *c_β_* their associated parameters [24].

### 4.4 HFS microrheology

As the HFS measurements work on suspended bodies, we simply transferred the spheroids from their culture well plates to Petri dishes. We used the HFS device in a configuration described previously[18, 66], using nozzles with 52-60 *µ*m diameter. We performed the measurements in PBS (Sigma-Aldrich, Poznań, Poland), at room temperature. To avoid effects arising from cell death, we performed the experiments in small sets of 4-5 spheroids, which would typically take 30-40 minutes to complete. Between each set of measurements, we washed the tip with a 10% solution of Helyzime (Braun, Germany), Isopropanol (Sigma-Aldrich, Poznań, Poland), and deionized water. Whenever we observed cell debris within the nozzle which could not be removed by washing, we sonicated the tip for about 20 seconds, in isopropanol and deionized water. We ran all the experiments in pressure control mode, with an aspiration pressure set point of 300 Pa and a suction rate of 50 Pa/s. We measured the dynamic response between 0.05 and 20 Hz, in 9 log-evenly spaced frequencies. We set a constant pressure oscillation amplitude of 50 Pa, waiting 2 seconds in between oscillations. We collected data for each spheroid on 4 well-separated locations, by releasing and recapturing them with the help of the pressure controller and a micromanipulator to reposition the suction tip. We calculated the complex modulus as we detailed a previous work [66], using the equation:

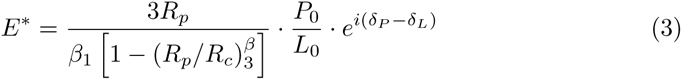

where *R_p_* is the pipette radius (between 52 and 60 *µ*m), *R_c_* is the sample radius, *β*_1_=2.0142, *β*_3_=2.1187, *P*_0_ and *L*_0_ are the amplitudes pressure and aspirated length respectively, and *δ_P_* and *δ_L_* their respective phase shifts. We corrected for hydrodynamic drag and analyzed the data as described in our previous work[18]. An example of the raw data is shown in the supplementary figure S7. After obtaining storage and loss moduli as a function of the frequency, we fit the DMA data to the Poynting-Thompson model in the complex plane. The equation of the model takes the form:

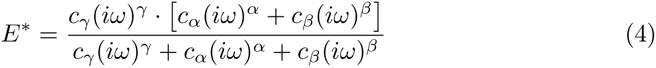

where, analogously to the AFM fit, *α*, *β*, *γ* are indices of derivation for three springpots, with their associated terms.

When performing the fit, we initialized the parameters with the values obtained modeling the AFM data. In order to verify the need for a fractional model, we defined a classic Burgers model, ran the optimization procedure, and visually compared the result to the output of the fractional model.

### 4.5 AFM cross-section analysis

After culturing the spheroids for 3 to 14 days, we mixed them with optimal cutting temperature compound (OCT, ThermoFisher), and froze them at -80 C. We cut the frozen samples in 20 *µ*m thick serial sections at -20 C using a cryostat (Leica Biosystems), and mounted them on poly-L-lysine coated slides. For the indentation experiments, we thawed the cryosections and washed the OCT in a PBS bath. We used NSC36 cantilevers (0.6 N/m, tip height 12-18 *µ*m and half-open angle of 20 degrees), and set approach velocity to 3 *µ*m/s and force setpoint to 30 nN. We measured the cryosections submerged in PBS, at room temperature.

Analogously to the microrheology data, we analyzed the force distance curve with the Hertz model, fitting the data up to 1.5 um of indentation using pyFM.

### 4.6 HFS tomography

To analyze the spheroids as layered bodies, we retrieved their depth resolved displacement maps. A crucial aspect to ensure repeatability was to automate the spheroid localization in the spectrometer trace. To do so, we first recorded the spectrum of background signal, in the same medium of the measurements. We subtracted the background from the experimental signal, resampled the result linearly in k-space, calculated the FFT with 2x zero padding, and normalized it. We then identified the pressure sensor as the local FFT intensity maximum located between 100 and 200 *µ*m from the reference fiber surface, and the front and back of the spheroids as the first and last points with intensities grater than 0.7 located past 300 *µ*m from the reference fiber respectively. We then demodulated the phase of the regions of interest and calculated the displacement along the spheroid diameter as:

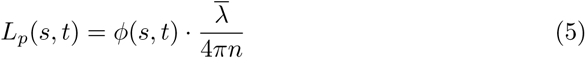

where *s* is the position along the spheroid diameter, starting from 0 at the surface, *t* is time, *λ* is the average wavelength of the source, and *n* is the estimated refractive index of the spheroid. We obtained *n* on a per line basis, averaging the optical cross section measurements for each spheroid and dividing them by the diameter measured in the brightfield image. We obtained values ranging from 1.37 to 1.42, in line with what has been previously reported in the literature [67].

We then downsampled the signal to 100 Hz to speed up data processing, and smoothed the spheroid field displacement data using a 2D Gaussian filter with 2×2 kernel size. As we limited the tomography analysis to elastic behavior, we extracted the displacement profile corresponding to a suction pressure of 200 Pa, as a compromise between obtaining a well developed displacement profile and the need to minimize viscous relaxation, to be able to relate the data to the AFM measurement on cryosections. As the spheroids were much larger than the pipette diameter, more than half the points and the distal end of the spheroids (away from the pipette nozzle) would measure the same displacement. This is because the strain in this region is zero, but as the sample is floating and the material is incompressible, this region moves rigidly. As these points do not contain useful information for the mechanical model, we truncated the arrays 120-150 *µ*m away from the surface, i.e. approximately 5 times the pipette radius. We saved two arrays, one containing the displacement data, and one position where the displacement was recorded.

We then passed the array to a modified version of ALVA[36], where we fitted a layered elastic model consisting of a two layers, optimizing the elastic moduli of the top and bottom layer, and the thickness of the top layer. We considered the pressure exerted by the nozzle on the spheroid, as it counteracts the suction pressure, by adding three distributed circular loads: one corresponding to the suction, and two corresponding to the pressure ring applied by the nozzle rim. The nozzle pressure was calculated with a simple force balance as:

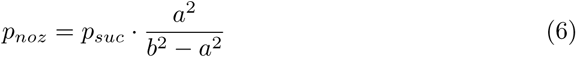

where *p_suc_* is the suction pressure, and *a* and *b* are the inner and outer nozzle radii respectively, measured from an SEM image of the nozzle. For the optimization, we constrained the top thickness to be at least one cell thick (8 *µ*m), and the elastic moduli to be between 10 Pa and 100 kPa. As ALVA does not account for rigid body displacement, we rigidly translated the curves by adding a constant term to the solution and treated the rigid body displacement as an optimization parameter (constrained within 3% of the experimentally measured value). We then filtered the curves based on their *R*^2^ value, keeping only those *>*0.95.

We validated the numerical approach in two ways. First, we used ALVA to generate displacement profiles for a two-layered system with different top thicknesses and E ratios, and then used this data as input for the optimization problem. In this case, we found the method to be robust and self-consistent, as ALVA was able to correctly retrieve both the magnitude of the moduli as well as the layer thickness, with zero error. We then validated this method against finite elements simulations, obtained as described in the Methods section above. As ALVA does not account for surface curvature, we verified the differences in displacement field associated to varying sample radii with a fixed pipette diameter. Given the range of displacements and minimum diameter, we found the error associated with the finite size to be less than 3%, in line with the results reported by a previous study [68]. Last, we used FEM data as input for ALVA. In this case, we found larger discrepancies, typically 8-10% for the top layer, 10-40% for the bottom layer, and about 20% for the thickness. The origin of these discrepancies is most likely to be found in the simplified boundary conditions we applied. Whilst the pipette nozzle exerts a balancing pressure on the sample, its distribution is not homogeneous over the contact region. A more accurate boundary condition would be a zero displacement in the z directions for the portion of the domain in contact with the nozzle. Comparing the ratio estimate by ALVA with the ground truth, we found ALVA to underestimate the top/mid ratio, meaning the claims about layered mechanical heterogeneity hold despite the estimation error. We found this simplification satisfactory considering the convergence error for the FE model and the fact that the underestimation was monotonically increasing with the ratio of elastic moduli.

### 4.7 Finite Element Modeling

We simulated a 2D axisymmetric layered sphere subject to suction using ANSYS APDL 2021-R2. We modeled the geometry using eight nodes elements with mixed u-P formulation. We considered the pipette surface to be rigid and frictionless, and all materials to isotropic and incompressible, and modeled the contact using the augmented Lagrangian formulation. We ran a convergence analysis and calculated the results shown in this paper for a model composed of approximately 30000 nodes, with Von Mises stress values converging to less than 5%.

### 4.8 Intensity Fluctuation Analysis

We implemented the LIV and OCDS metrics as reported by a previous study [45]. Briefly, we calculated LIV as:

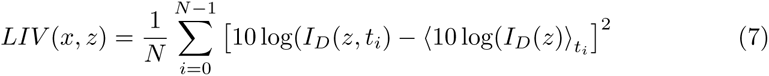

where *N* is the number of frames (800 in this case), *I_D_* is the signal intensity as function of time *t*and depth *z*, and the ⟨⟩ brackets indicate the time average. We calculated OCDS by first computing the autocorrelation signal between 10 and 400 ms as:

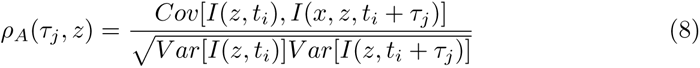

where *Cov* and *V ar* are the covariance and variance of the signal respectively, and *τ_j_* indicates successive time lags. We then calculate the slope of the decay as a function of delay time, and took its absolute value as OCDS.

### 4.9 Histology

We collected the spheroids cultured for 3 and 14 days in 1.5 ml Eppendorf tubes. We fixed the samples with 10% neutral buffered formalin (Sigma-Aldrich, St. Louis, MO. USA) for 15 minutes. Slides were dehydrated in a series of diluted alcohol solutions: 50%, 80%, 96%, and 100%. Each dehydration step lasted 15 minutes. We then treated the samples with Xylene (Sigma-Aldrich, St. Louis, MO. USA), then removed it and embedded the spheroids in paraffin blocks. We cut 6 *µ*m thick slices using a microtome (Biocut 2035, Leica Instruments GmbH), and mounted them on poly-L-lysine coated slides (Menzel-Glaser, Thermo Scientific).

Deparaffinized sections were stained with hematoxylin and eosin, as well as using histochemical methods: Masson’s trichrome to distinguish cells from surrounding connective tissue and picrosirius red stain for collagen components (collagen rich ECM) to check their possible correlations with nanomechanical and rheological changes of bladder spheroids.

### 4.10 SDS-PAGE

After three days of incubation, we collected the spheroids, washed them in PBS, and added 10 *µ*l of solubilizer, prepared by dissolving in water 1M TRIS-HCL pH 6.8, Sodium Dodecyl Sulfate (SDS), glycerol and 2-Mercaptoethanol. We first heated all the solutions containing the spheroids to 95° C for 5 minutes, then sonicated them by using an ultrasonic homogenizer (BioLogics Model 300 V/T) three times at intervals of 5-10 seconds. We used a fluorimeter (Invitrogen Qubit, Thermo Fisher Scientific) to record theprotein concentration in the lysate. For each sample, 20 mg of protein was loaded in polyacrylamide gel for electrophoresis (4-15 % Mini-PROTEAN Gels, BioRad) and run in electrode buffer (10x Tris/Glycine/SDS, BioRad). The electrophoresis was conducted at a voltage of 60 V for 20 minutes and then at 120 V for two hours.

### 4.11 Western Blot and Immunodecoration

After the SDS-PAGE, we transferred the material to a PVDF blotting membrane using a transfer buffer (10x Tris/Glycine Buffer for Western Blots, BioRad). We performed a wet transfer at 4°C overnight, under a constant electric current of 15 mA. To prevent non-specific background bindings, we first incubated the membrane in a blocking solution containing 3% of Bovine Serum Albumin (BSA) solution for 3 hours at 4°C under gentle agitation and then washed it three times in TBS/Tween20 (TBST) buffer (Sigma Aldrich). Then, we incubated the membrane with the primary antibodies N-cadherin (mouse monoclonal [8C11] to N-Cadherin, ThermoFisher) and E-cadherin (E-cadherin Monoclonal Mouse clone NCH-38, DAKO), in blocking solution (TBST) overnight at room temperature and washed again in TBST buffer. We verified the presence of the protein bands by using a secondary antibody (anti-Mouse IgG (whole molecule)-Alkaline Phosphatase antibody).

### 4.12 Confocal Imaging

We used the following protocol to stain spheroids at both 3 and 14 days: we collected the samples in 1.5 ml Eppendorf tubes and fixed them in 3.7% PFA for an hour. The samples were then treated with 1% cold Triton X-100 overnight at 4°C, washed three times in PBS for two minutes each, and then washed once more in PBS. Then, we added 1% cold BSA solution, for 3 hours at 4 C. Once again, we washed the samples in PBS. Next, we incubated the spheroids overnight at 4 C with phalloidin conjugated with Alexa Fluor 488 (1:20, in PBS). The following day, we substituted the dye with Hoechst 33342, diluted 1:5 in PBS, and incubated in the same way. The following day, we washed the samples in PBS, moved them to an 18 well, glass bottom slide (Ibidi), with the addition of an anti-shading solution (Thermo Fisher) with the same refractive index as the oil used for the immersion objective (1.52). We recorded the images using a Leica TCS SP8 WLL confocal microscope, using a 63x objective lens (HC PL APO CS2, NA 1.40).

### 4.13 Brightfield Image Analysis

Using an inverted microscope with phase contrast illumination, we captured pictures of the spheroids over the course of the two-week culture to track the spheroid growth and morphological changes. To obtain the projected area and circularity metrics, we segmented the spheroid images using Segment Anything Model[69]. We then used the binary masks to calculate area *A* and perimeter 2*p* using the regionprops function in Matlab R2022b. Circularity was defined as:

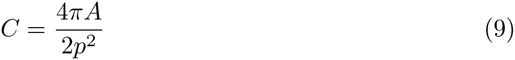

To obtain the degree of spheroid fusion, we fit a circle to each spheroid, and calculated the angle *θ* between the line connecting the centers of the circles and the linne passing through one of the circles and one of the intersection points. The degree of fusion was calculated as *cos*^2^*θ*.

### 4.14 Quantitative Histology

We performed quantitative analysis of the H&E staining using a pre-trained convolutional neural network (Stardist 2D, Versatile H&E model[70]), accessed through ImageJ. An example of the segmentation result is available in the supplementary figure S8. We extracted coordinate list, area, perimeter, aspect ratio, circularity for each of the nuclei. We imported the results in Matlab R2022b for further processing. We iteratively selected the outermost set of nuclei, and recorded their average feature, as well as the distance from the initial set boundary. We then grouped the nuclei in three sets, under the assumption that the first 25 *µ*m contain mostly proliferating cells, and that beyond 100 *µ*m nutrients have lower concentration. In practice, as one would expect, both the aspect ratio and area vary continuously, both in the radial and tangential directions (see Supplementary figure S9). That being said, a dependence of AR as function of radial position is apparent in all lines. We repeated the procedure on two slides per each condition, and found the results to be very consistent. We calculated the 3 to 14 days compaction as:

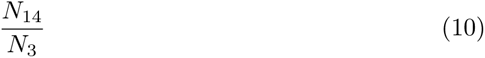

where N is the nuclei density per *µ*m^2^ at 14 and 3 days, as denoted by the subscripts. We estimated the area of each region using the boundary definition described above. We compared the nuclei aspect ratio in between regions for each spheroid, as a proxy for cell shape, using a Mann-Whitney U test.

### 4.15 Statistics and Reproducibility

In all mechanical characterization experiments, measurements were conducted on a minimum of 10 spheroids. All statistical tests were conducted using a Mann-Whitney U test, assuming a p value lower than 0.05 could be considered statistically significant.

## Supplementary information

This article has supplementary information.

## Code Availability

All the code used to analyze the data is freely avalible at: https://github.com/max-bera/HFS_analysis and https://github.com/asmusskar/ALVA

## Author Contributions

K.G.: Conceptualization, Methodology, Investigation, Formal Analysis, Visualization, Writing - Original Draft

M.B.: Conceptualization, Methodology, Investigation, Hardware, Software, Formal Analysis, Visualization, Writing - Original Draft

A.S.: Software, Formal Analysis, Writing - Review and Editing

G.P.F.: Investigation, Formal Analysis, Writing - Review and Editing

J.P.: Investigation, Formal Analysis, Writing - Review and Editing

J.L.A.: Software, Formal Analysis, Writing - Reviewing and Editing

I.B.A.: Writing - Reviewing and Editing

M.L.: Funding acquisition, Writing - Reviewing and Editing

## Acknowledgments

This work was financially supported by the H2020 European Research and Innovation Programme under the Marie Sk-lodowska-Curie grant agreement “Phys2BioMed” contract no. 812772.

The authors express their gratitude towards Giulia Pilia, for the useful discussions, towards Laura Martinez Vidal, for the help in preparing the cryosections, and towards Ekrem Sahin, for the help in cross-validating the FE model.

