## Supplementary materials for "Multiscale Rheology of Aging Cancer Spheroids"

### RESULTS

1. **Extended images of Western Blot results obtained for 3-days-old spheroids.**

A)

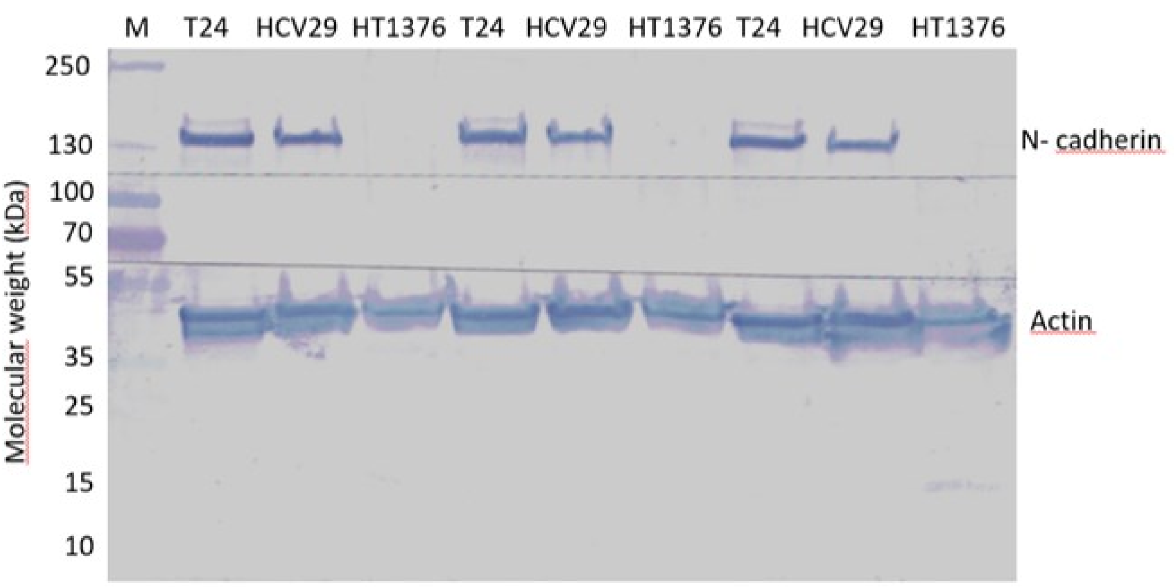


B)

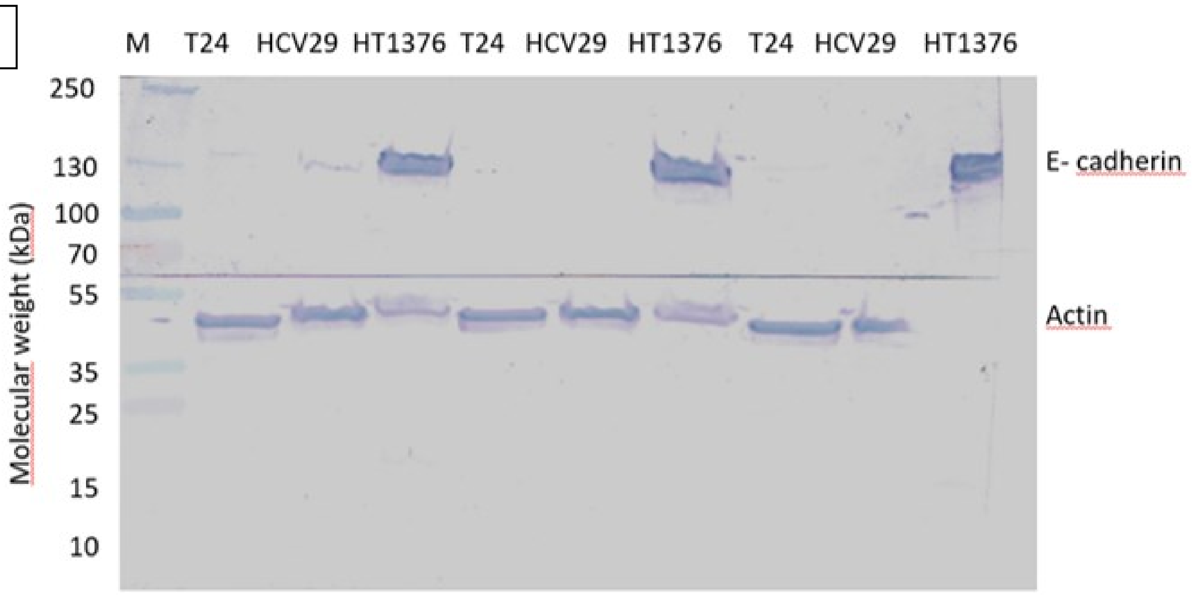


Supplementary Figure S1. Exemplary Western blots showing the expression of cadherins in bladder cancer spheroids. A) N-cadherin expression. B) E-cadherin expression. HCV29 and T24 spheroids express only N-cadherin, while HT1376 spheroids express only E-cadherin. M = protein marker, on the left the molecular weight of the reference bands (kDa) are reported, on the top of each lane the name of the cell lines is indicated.

2. **Extended version of the histological images of the spheroids used in figure 4.**

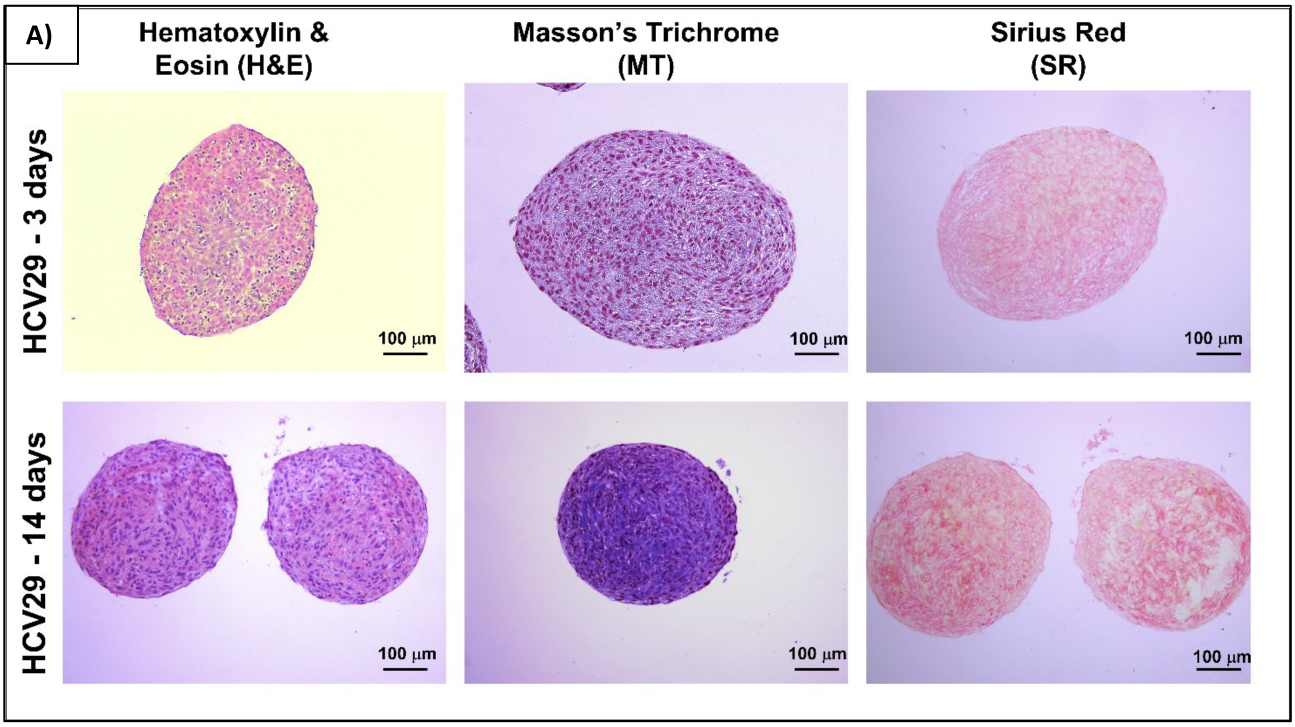


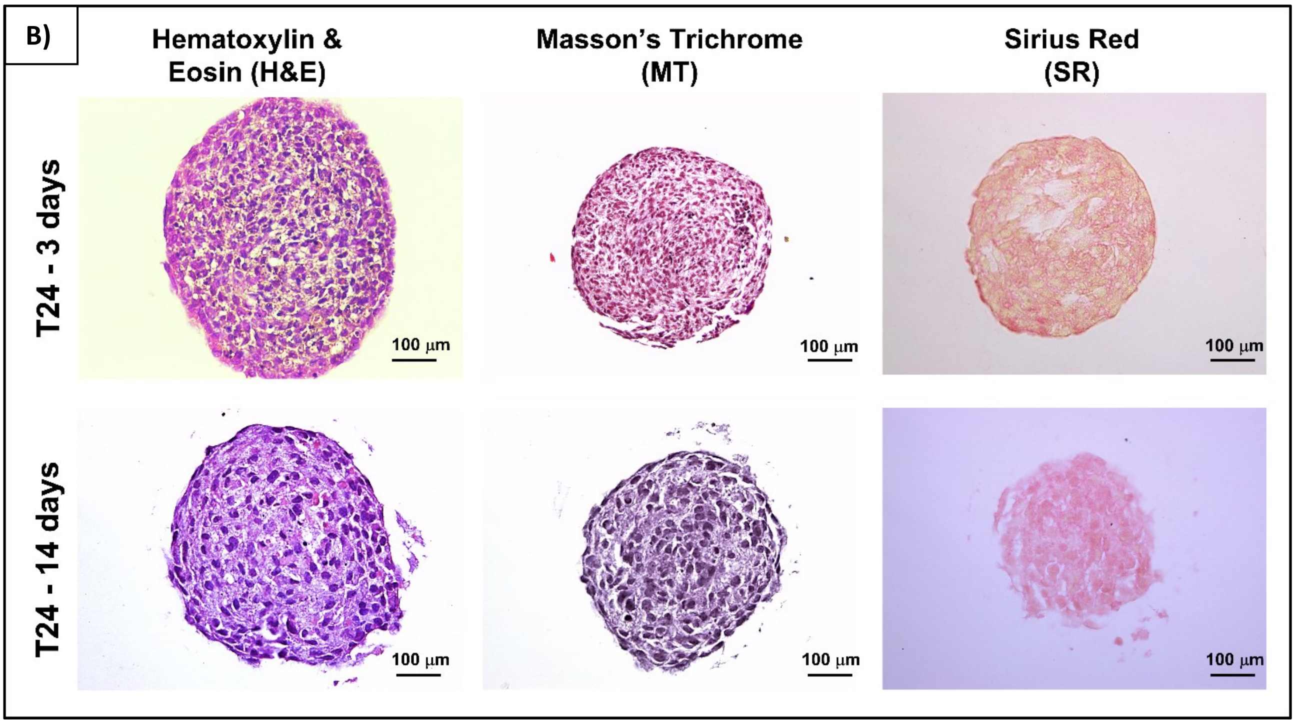

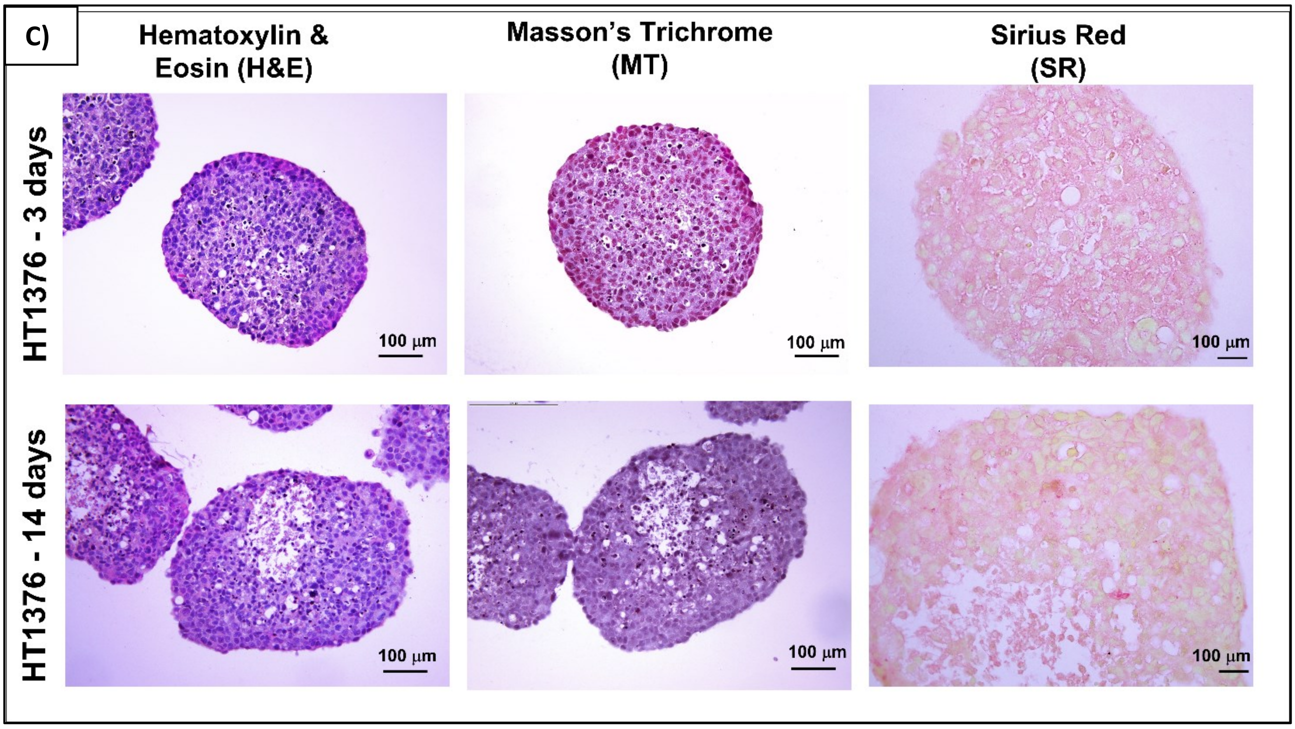


Supplementary Figure S2. Histological staining of the spheroids formed by HCV29 cells (A), T24 cells (B) and HT1376 cells (C) after 3 days and 14 days of culture. The spheroid sections were stained with Hematoxylin and eosin (H&E), Masson's Trichrome (MT) and Sirius Red (SR). H&E stains the cell nuclei in purplish blues by separating it from all the other structures, MT distinguishes cells from the surrounding connective tissue stained in blue, and SR stains in red the collagen present in the extracellular matrix (ECM).

3. **Extended version of the confocal images showing the organization of actin cytoskeleton inside spheroids in figure 4.**

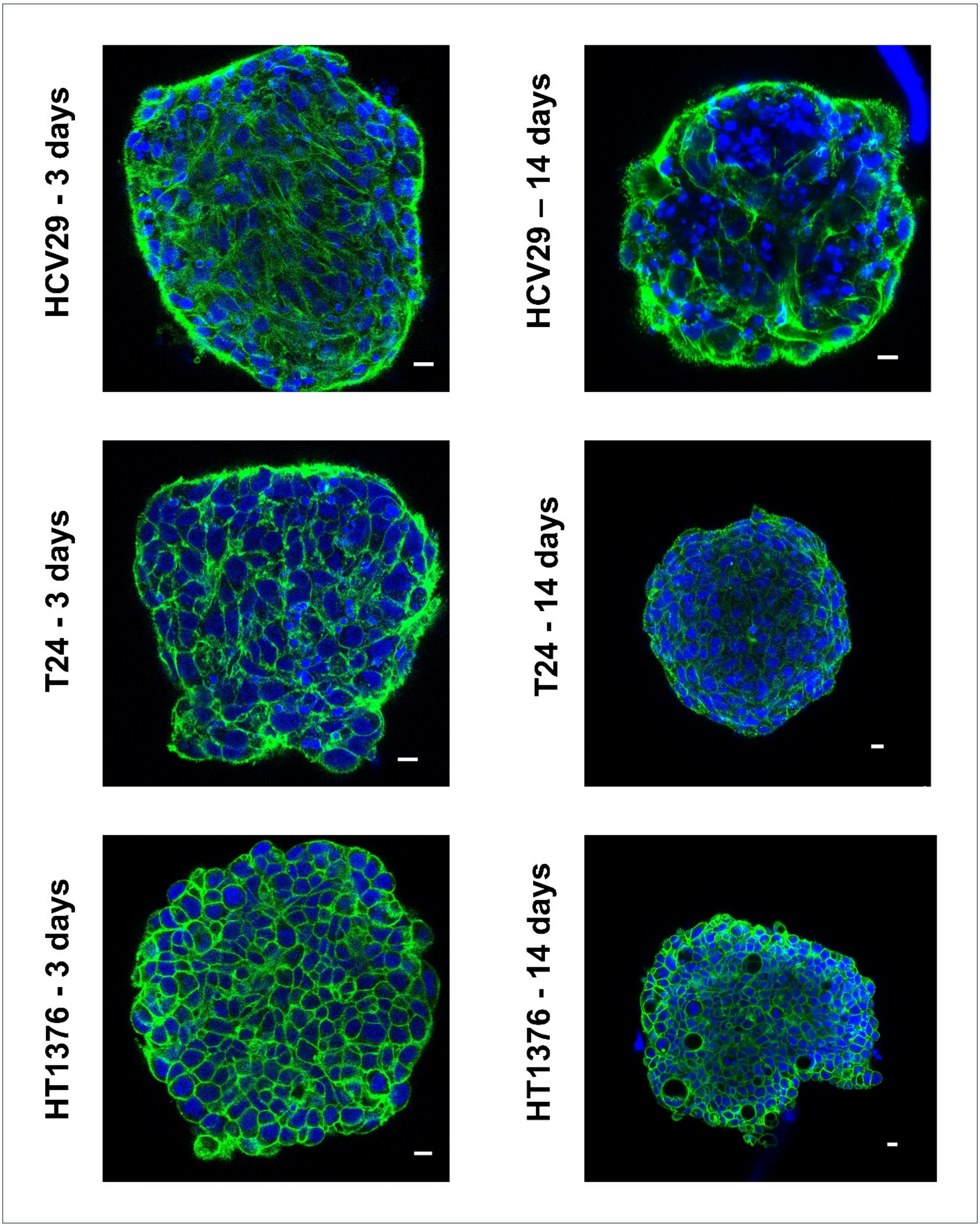


Supplementary Figure S3. Confocal images showing the organization of actin cytoskeleton inside the spheroids. Green: F-actin, stained with phalloidin conjugated with Alexa Fluor 488, blue: nuclei, stained with Hoechst 33342. Scalebar = 10 μm.

**4. Fluorescence images of spheroid cryosections that were measured with the AFM.**

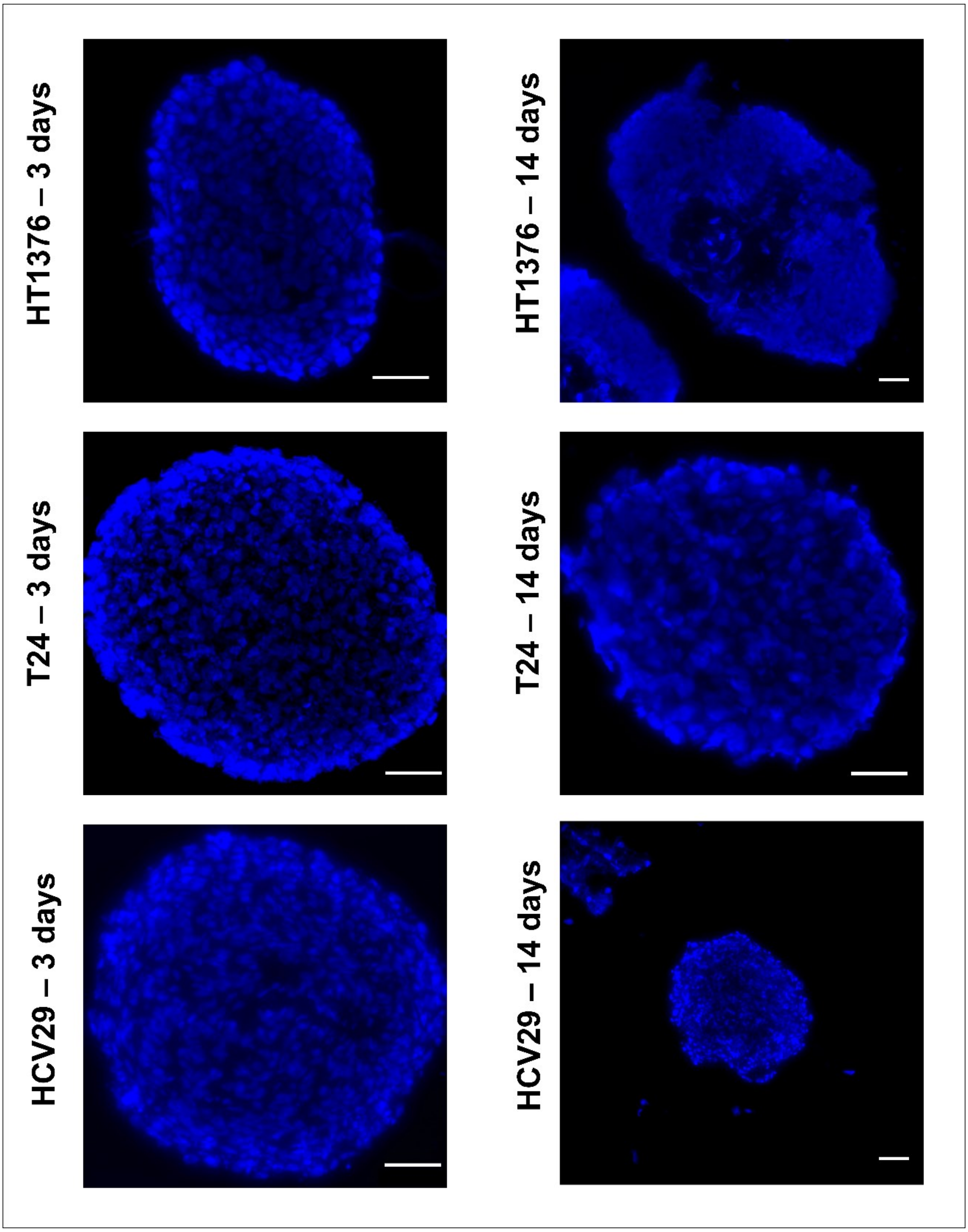


Supplementary Figure S4. Fluorescence images of the cryosections of the spheroids showing the nuclei, stained with Hoechst 33342. Scalebar = 50 μm.

**5. Results of the Modified Maxwell Fit on HFS-DMA data.**

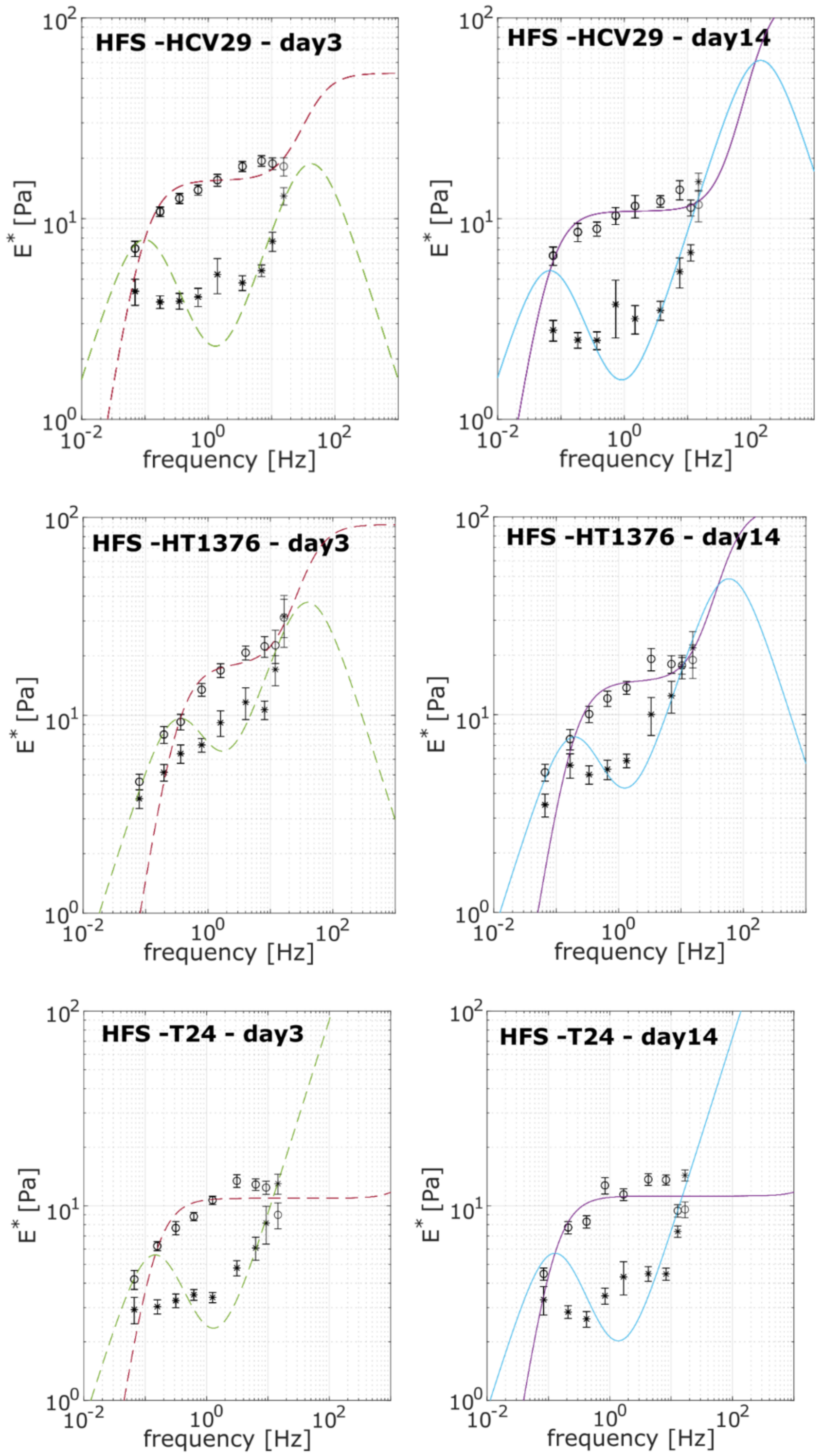


Supplementary Figure S5. Results from HFS-DMA data obtained after Modified Maxwell fit. From visual inspection, it is evident that this topological configuration of springs and dashpot cannot adequately capture the full range in which the spheroid behaves as a solid.

### METHODS

#### 1. Exemplary force curve from AFM microrheological measurements.

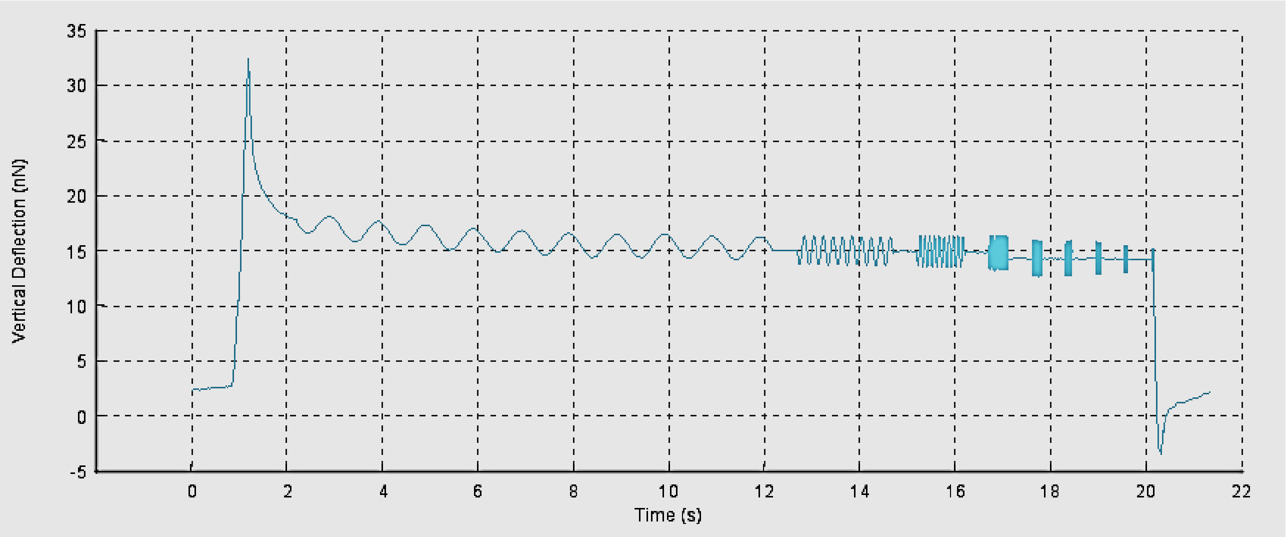


Supplementary Figure S6. Exemplary force curve obtained from the AFM microrheological measurements. The dynamic response was set between 1 and 250 Hz in 9 log-evenly spaced frequencies. The following parameters were set: a constant oscillation amplitude of 50 nm for 10 oscillations, sampling 600 points per period, and waiting 0.5 seconds between test frequencies.

#### 2. Determination of shear storage and loss moduli from AFM microrheological measurements.

Rheological properties of cells measured by AFM relied on the theoretical approach developed by Alcaraz et al. in 2003 (Alcaraz et al., 2003). For small oscillation amplitudes, the complex shear modulus E* for pyramidal indenter with a half open-angle $\Theta$ is calculated and can be described by the following equation:

$$E^* = \frac{2\,(1-\nu^2)}{3\,\delta_o\,\tan\theta}\;\frac{F\,(\omega)}{\delta\,(\omega)} \qquad (3)$$

where $F$ is the load force and $\delta$ is the indentation depth. The Fourier transform of the force and indentation at the frequency $\omega$ can be written as:

$$F(\omega) = F_A \cdot e^{i\varphi_F}$$

$$\delta(\omega) = \delta_A \cdot e^{i\varphi_\delta}$$

$F_A$ and $\delta_A$ are the amplitudes of the force and indentation at the angular frequency $\omega$ (= $2\pi{\cdot}f$) after the Fourier transforms, respectively.

Phase shift $\Delta\varphi$ between the force and deformation amplitude is $\Delta\varphi = \varphi_F - \varphi_\delta$

Then,

$$F(\omega) = F_A \cdot (\cos\Delta\varphi + i \cdot sin\Delta\varphi)$$

$$\delta(\omega) = \delta_A$$

Finally, the equation (3) can be re-written as:

$$E^* = E' + iE" = \frac{2\,(1-\nu^2)}{3\,\delta_o\ tan\,\theta}\ \frac{F_A \cdot (cos\,\Delta\varphi + isin\Delta\varphi)}{\delta_A}$$

By the relationship between the elastic and shear moduli, i.e., $G = \frac{E}{2\,(1-\nu)}$, the shear storage $G'$ and loss $G''$ moduli are:

$$G'(\omega) = \frac{1-\nu}{3\,\delta_o \tan\theta}\ \frac{F_A}{\delta_A}\ \cos\Delta\varphi \quad (4)$$

$$G^{"}(\omega) = \frac{1-\nu}{3\,\delta_o \tan\theta}\ \frac{F_A}{\delta_A}\ \sin\Delta\varphi \quad (5)$$

where *G'(ω)* measures the elastic energy that is stored and recovered during the oscillation, while *G''(ω)* stands for the loss modulus that considers the energy dissipated.

[Information obtained from the JPK Manual]

#### 3. Example of dynamic mechanical analysis experimental data from HFS.

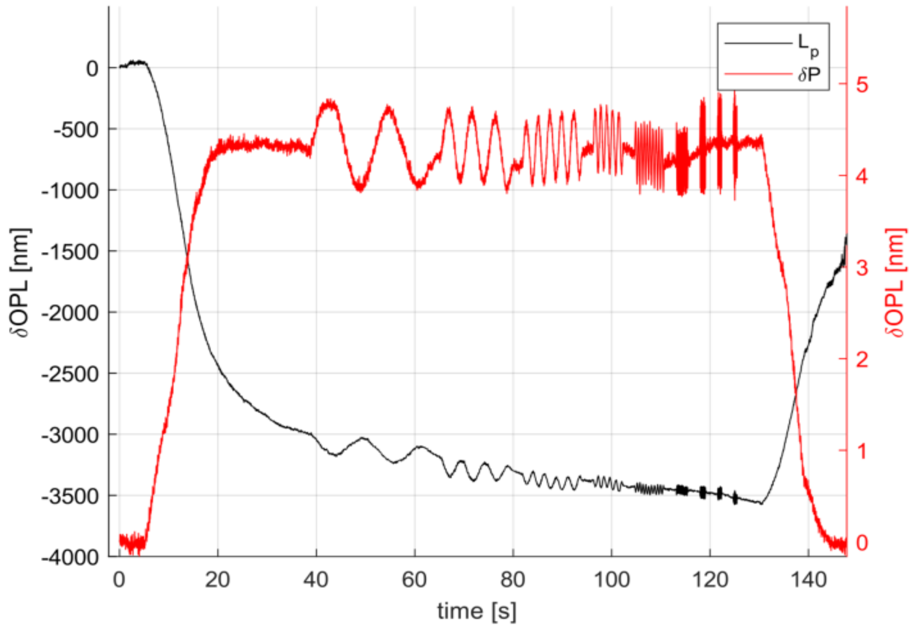


Supplementary Figure S7. Exemplary force curve obtained from the HFS DMA measurements. The dynamic response was measured between 0.05 and 20 Hz, in 9 log-evenly spaced frequencies. The pressure oscillation amplitude was set constant at 50 Pa, waiting 2 seconds in between each oscillation.

#### 4. Example of nuclei detection using Stardist.

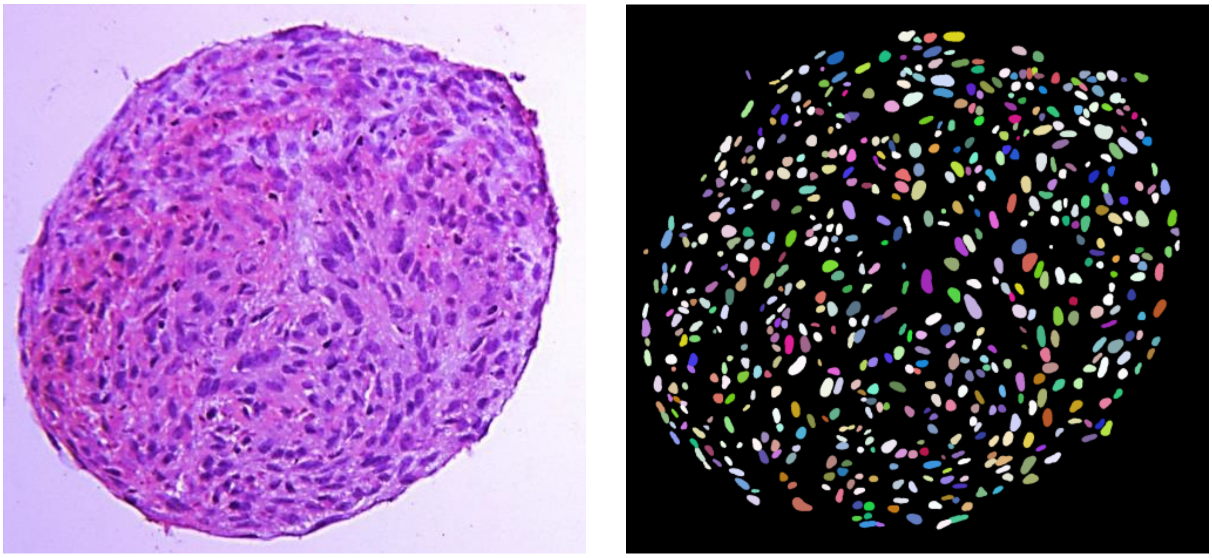

Supplementary Figure S8. Example of a segmentation result obtained from quantitative analysis of H&E staining using a pre-trained convolutional neural network (Stardist 2D, Versatile H&E model) with ImageJ software.

**5. Example of the aspect ratio (AR) and the area of the nuclei.**

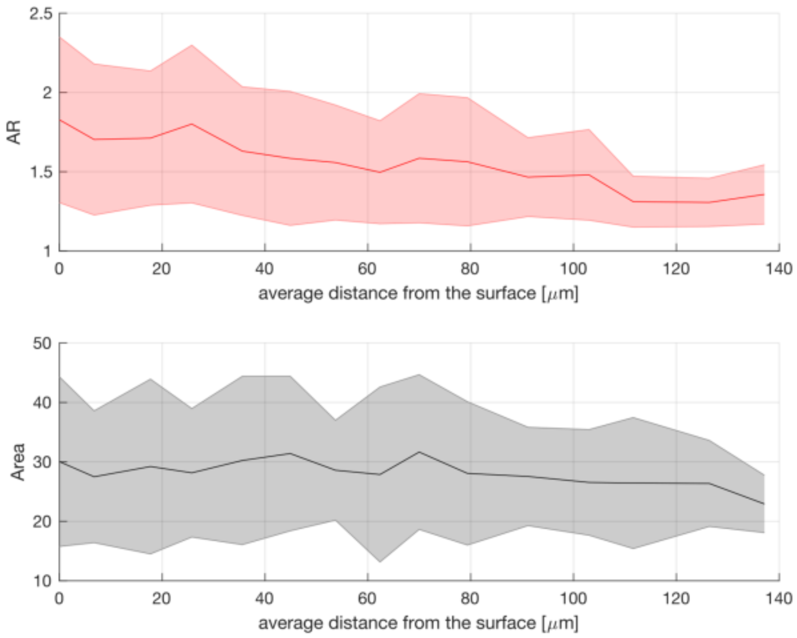


Supplementary Figure S9. Example of the distribution of the aspect ratio (AR) and the area of the nuclei extracted from the segmentation analysis. The results are shown in relation to the average distance from the surface, assuming that the first 25 μm is composed of proliferating cells, and that beyond 100 μm there is a decay of the nutrients concentration.
